# Lake ecosystem responses to runoff variability across time and space

**DOI:** 10.64898/2026.06.18.729811

**Authors:** Silke Langenheder, Jorrit P. Mesman, Nils Kreuter, Dolly Kothawala, Gabriela Ágreda-López, Akif Ari, Stella A Berger, Susana Bernal, Bence Buttyán, Berenike Bick, Nuria Carabal, Nuria Catalan, Eleni A Charmpila, William Colom Montero, Pelin Ertürk Arı, Inge Elfferich, Johanna Exner, Bence Gergácz, Emma Gray, Anika Happe, Congcong Jiao, Kevin Jones, Nusret Karakaya, Antonija Kulaš, Anna Lupon, Clara Mangold, Clara Mendoza-Lera, Jens C. Nejstgaard, Jimmy C. Oppong, Angela Pedregal-Montes, Nuria Perujo, Juha Rankinen, Tobias Rütting, Johanna Sjöstedt, Maren Striebel, Katerina Symiakaki, Eline van Dam, Simon Wentritt, Majd M Yaqoob, Kadir Yıldız, Ingrid Sassenhagen

## Abstract

Inland waters in the Northern Hemisphere are experiencing increased annual runoff due to higher overall precipitation as well as intensified short-term events such as heavy rainfall, floods and storms. These events affect the total loading and variability of inputs of allochthonous, coloured dissolved organic matter (cDOM) and inorganic nutrients into lakes. Previous studies have shown that increased total cDOM and inorganic nutrient loads affect phytoplankton biomass and metabolic rates, but it is unknown how the effects of different cDOM and nutrient pulse scenarios are modified by spatial and seasonal differences in lake characteristics. Here, we conducted a coordinated, standardized mesocosm experiment across three lakes with different ambient cDOM and nutrient concentrations. In two of these lakes, the experiment was implemented in two seasons. The same total amounts of cDOM, nitrate and phosphate were added to all mesocosms, but in pulses that differed in intensity and frequency. We found that pulse intensity and frequency affected chlorophyll *a* and phycocyanin concentrations and metabolic rates, i.e. gross primary production and respiration, differently. Specifically, more pronounced effects were found in response to the extreme pulse scenario compared to those with more frequent, smaller pulse additions. Furthermore, the effects were mainly temporary and varied more among lakes than between seasons. The clearest differences between the extreme and more gradual runoff scenarios were found in the lake with the lowest background cDOM and nitrate concentrations, likely because lower light limitation and possibly stronger initial N-limitation caused a faster response to the nutrient addition. Our results highlight that both antecedent lake conditions and characteristics of runoff events can affect phytoplankton biomass and metabolic rates and that comparative experimental approaches are needed to reveal the complexity of the responses.

## Introduction

Ecosystems are increasingly affected by gradual and abrupt, unpredictable changes in environmental factors induced by climate change related to extreme weather events (O’Gorman and Schneider 2009, Zhang et al. 2021, IPCC 2023). So far, global change research has largely focused on the effects of long-term alterations in mean environmental conditions even though extreme, short-term events may be equally or even be more important for ecological communities and ecosystem processes and influence both short- and long-term ecosystem responses (Jentsch et al. 2007, Jennings et al. 2012, Forsman et al. 2016). Inland waters, for example, offer a clear example of such vulnerability to long-term gradual as well as short-term extremes as they are already experiencing increased annual runoff due to higher precipitation as well as more variable and intense short-term events such as rainfall, floods and storms (Madsen et al. 2014, Stockwell et al. 2020, Woolway et al. 2020). These changes affect the total loading and variability of pulse events of allochthonous, coloured dissolved organic material (cDOM) and inorganic nutrients inputs from the surrounding catchment into lakes (Jennings et al. 2012, Zwart et al. 2017, Stockwell et al. 2020). External loading changes underwater light conditions due to browning, elevates inorganic nutrients, such as nitrogen (N) and phosphorus (P) and dissolved organic carbon (DOC) concentrations and causes shifts in carbon and nutrient stoichiometry (e.g. Klimaszyk et al. 2015, Corman et al. 2023).

Inputs of cDOM can potentially affect ecosystem responses, including primary production, phytoplankton growth dynamics and bacterial production, and consequently alter key metabolic processes such as gross primary production (GPP), ecosystem respiration (ER) and net ecosystem production (NEP = GPP - ER) (Hanson et al. 2003, Jeppesen et al. 2021). These alterations can further cause shifts in food web structure, energy fluxes to higher trophic levels, and greenhouse gas emissions (Hanson et al. 2003, Cole et al. 2007, Tranvik et al. 2009). cDOM inputs can lead to increases in ER due to bacterial utilization of allochthonous DOC (Tranvik et al. 2009, Zwart et al. 2016) while the response of primary producers to runoff inputs is more variable and difficult to predict (Solomon et al. 2023). Increases in inorganic nutrient concentrations can promote increases in phytoplankton biomass and GPP (Deininger et al. 2017, Jeppesen et al. 2021), whereas high cDOM inputs can have the opposite effect because it increases light limitation for algae (Sadro and Melack 2012, Seekell et al. 2015, Deininger et al. 2017, Bergström and Karlsson 2019, Olson et al. 2020, Isles et al. 2021). This trade-off between nutrient and light availability can result in hump-shaped relationships between cDOM concentrations and GPP relationships (Kelly et al. 2018, Olson et al. 2020). Moreover, changes in nutrient availability and light conditions associated with runoff inputs can alter phytoplankton community composition, particularly the abundance, species composition and toxin production of cyanobacteria, which is of major environmental concern (Reichwaldt and Ghadouani 2012, Luimstra et al. 2020, Lyche Solheim et al. 2024).

In general, the effects of runoff loading on phytoplankton biomass and the different ecosystem metabolic rates can be highly variable across lakes and seasons due to differences in both local lake as well as runoff characteristics. For example, ecosystem responses to runoff inputs can be mediated by interactions with foundation species (Lürig et al. 2021), fish presence (Cottingham and Schindler 2000), antecedent lake conditions (e.g. turbidity, abiotic nutrient and light conditions) (Thayne et al. 2022) and load N:P ratios (Corman et al. 2023). Seasonal differences related to light and nutrient uptake rates of phytoplankton could also be important. For example, it has been shown that phytoplankton communities are dominated by species with high nutrient uptake affinities when nutrient concentrations are low, which could lead to more rapid responses to external nutrient inputs (Pranger et al. 2025). Further, ecosystem responses can vary depending on the magnitude of cDOM and nutrient loading during runoff events (Jennings et al. 2012, Meng et al. 2017, Kelly et al. 2018, Corman et al. 2023). Responses might, however, not only depend on characteristics of individual runoff events but also on antecedent climatic conditions and the sequence and characteristics of previous events, leading to legacy effects.

Global change experiments aim to draw general conclusions from case studies, but often fail to do so because results from ecological experiments are highly context dependent (De Boeck et al. 2015, Catford et al. 2022). Thus, results may be influenced by the local setting of each study, often leading to contrasting responses to common drivers of environmental change (Mahdy et al. 2015, Urrutia-Cordero et al. 2021a). Spatially replicated experiments that study responses to standardized manipulations with the same equipment and methods are, therefore, crucial tools to overcome this limitation (De Boeck et al. 2015, Urrutia-Cordero et al. 2021b). In addition to spatial influences, it is important to address how temporal differences in environmental conditions and community composition within systems affect the outcome of experiments with standardized treatments. Examples from “single system experiments” replicated in time could, for example, show strong year or season-specific effects in response to the same manipulation (Nejstgaard et al. 2006, Beisner and Peres-Neto 2009, Werner et al. 2020).

Here, we conducted coordinated, controlled mesocosm experiments that were replicated in space (three lakes) and time (two seasons in two of the lakes) to investigate how different cDOM, N, and P loading scenarios affect phytoplankton biomass and lake metabolic rates, which we measured using high-frequency measurements of chlorophyll *a* (chl-*a*), phycocyanin and dissolved oxygen concentrations. Specifically, we manipulated the number, intensity and frequency of cDOM and inorganic nutrient pulse additions during a 20-day ‘simulated rainfall’ period, followed by a ‘recovery’ period of 17 days without any further manipulations, to test whether ecosystem responses to differences in runoff persisted over time.

The total amount of added cDOM and inorganic nutrients (N and P) was kept the same between treatments and thus we applied fewer pulses of higher intensity on the one hand and more frequent but smaller pulses on the other hand, thereby creating a gradient of three scenarios ranging from ‘Extreme’ to more continuous ‘Daily’ cDOM and nutrient loadings. A control treatment without any additions was included as well. The experiments were replicated in space and time to compare effects among three lakes differing in water colour and nutrient concentrations. In two of the lakes, the experiment was also replicated in time (summer and spring) to compare whether ecosystem responses were related to seasonal differences in nutrient availability and plankton composition. This comparative approach is the first to investigate how chlorophyll *a* (chl-*a*) and phycocyanin concentrations indicative of phototrophically produced biomass (both phytoplankton and cyanobacteria) and ecosystem metabolic rates (GPP, ER and NEP) respond to different cDOM and inorganic nutrient pulse scenarios across temporal and spatial scales. Our aim was to test the following hypotheses:

(1) The three experimentally simulated runoff scenarios will result in different temporal patterns of chl-*a* and phycocyanin concentrations and metabolic rates due to differences in light and nutrient conditions in the water column over the course of the experiment. Specifically, based on previous results obtained from one single lake experiment that focused on changes in particulate organic carbon concentrations and oxygen saturation (Happe et al. 2025) we predict that extreme, abrupt pulses will at least temporarily lead to stronger increases compared to smaller, continuous additions.
(2) Responses of phytoplankton and metabolic rates to runoff scenarios will differ in lakes with different background cDOM concentrations. In humic lakes, overall weaker responses of GPP compared to ER are expected, resulting in lower NEP.
(3) Responses of phytoplankton and associated metabolic rates to runoff scenarios will differ between seasons due to differences in plankton composition and nutrient concentrations. Specifically, we predict stronger responses compared to background conditions in spring when inorganic nutrient concentrations are lower.

## Methods

### Experimental set-up

We conducted a total of five standardized *in situ* mesocosm experiments in the summer of 2022 and spring of 2023 at three field stations in Sweden (Bolmen Research Station, the Erken Laboratory and Skogaryd Research Catchment). They are part of the SITES AquaNet mesocosm infrastructures in three lakes (Bolmen, Erken, and Ersjön), respectively (Urrutia-Cordero et al. 2021b). We refer to each lake site by the station name (Bolmen, Erken and Skogaryd) throughout the study to be consistent with previous publications. A detailed description of the experimental setup can be found in the protocol collection in Langenheder et al. (2024). Briefly, mesocosms (700 L polyethylene containers, 1.2 m depth) were fitted into the openings of a Jetfloat platform and filled with approximately 550 L unfiltered lake water from the surroundings of the floating platforms from a depth of ca. 0.3 - 1 m (depending on the lake) using a pump (Meec tools 735-018, JULA AB, Sweden). The experiment started the following day (day 1) when the first sampling took place prior to the first cDOM and inorganic nutrient additions. The same total amount of cDOM and inorganic nutrients (N and P) was added to three runoff treatments, but at different frequency and magnitude: the ‘Extreme’ treatment consisted of a single large pulse addition (100% of the total amount), the ‘Intermittent’ treatment comprised seven pulses of variable intensity (5-30% of the total amount), and the ‘Daily’ treatment was achieved by daily additions (for 20 days) of low intensity (5% of the total amount) (Fig. 1). Finally, there was a Control treatment without any additions. Each treatment was replicated four times at each mesocosm site, resulting in a total of 16 mesocosms per site. cDOM and inorganic nutrients were added at the same time every day after all measurements, sampling and sensor cleaning were completed (see below). After 20 days, cDOM and inorganic nutrient additions were stopped and in order to follow up on a potential recovery phase, responses were monitored for another 17 days, for a total of 37 days per experiment.

**Figure 1.**
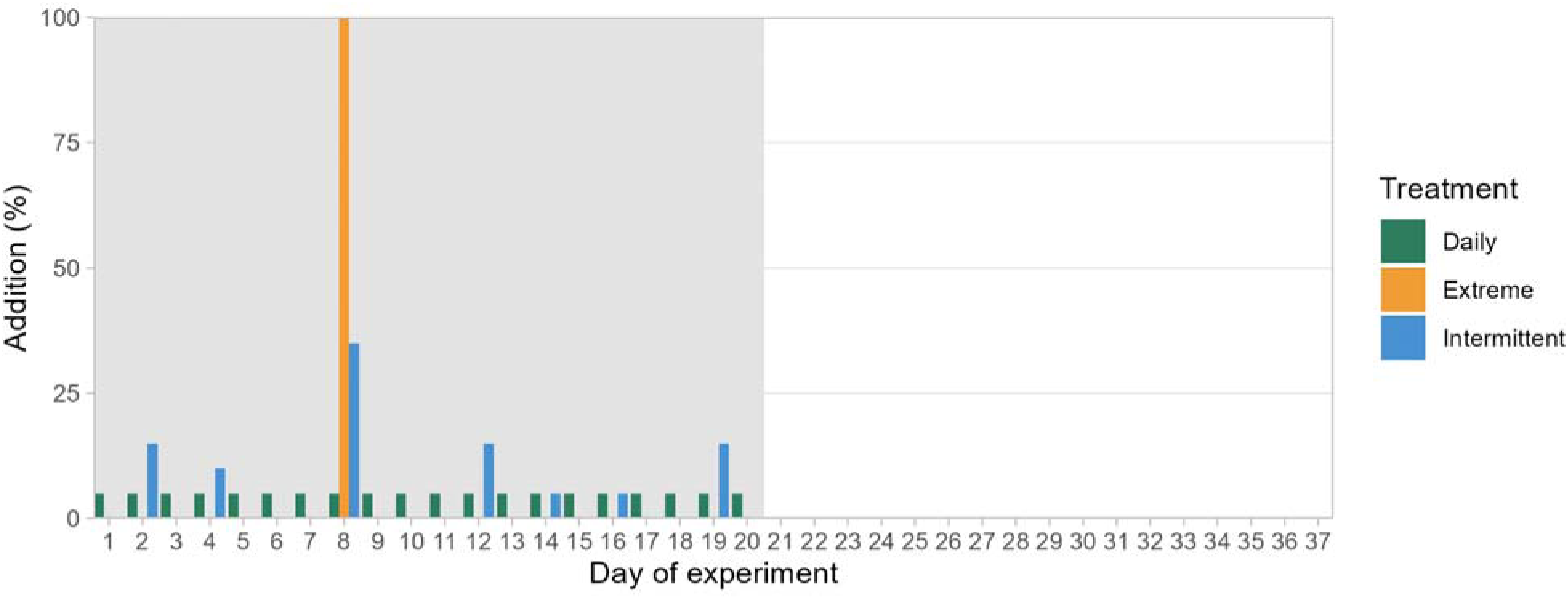
The three runoff scenarios, including 20 days with cDOM and nutrient additions: extreme, intermittent and daily, over the experimental study period of 37 days, followed by a 17 days long recovery period. The shaded area in grey is the simulated runoff period in which nutrients were added. The y-axis shows the fraction of total cDOM/nutrients (2 mg L^-1^ DOC, approximately, 50 μg L^-1^ P and 500 μg L^-1^ N (see methods section and Langenheder et al. (2024)).

Experiments conducted in summer 2022 in Erken and Bolmen were fully synchronized and ran from July 7 to August 12. Experiments were then run again in spring 2023 and at this point a third site (field station Skogaryd at lake Ersjön) was added to include an additional lake type. The start of the spring 2023 experiments differed slightly between the lakes (Bolmen: April 25, and Skogaryd: April 29, Erken: May 3) because of practical reasons (i.e. seasonal differences in the ice-off; see Langenheder et al. (2024)). The three lakes differ in background environmental conditions, including differences in water colour and trophic state (Urrutia-Cordero et al. 2021a, Urrutia-Cordero et al. 2021b). While DOC concentrations were similar in Erken and Bolmen, the two lakes differed clearly in water colour, with Erken being a clearwater lake and Bolmen a humic lake, whereas Skogaryd had the highest DOC concentrations and darkest water colour of all three lakes (Table S1). There were also clear differences in average daily photosynthetically active radiation (PAR) at 0.4 meters depth between the three lakes, with PAR decreasing from Erken to Bolmen and Skogaryd (Table S1). Further, there were differences in conductivity and pH (higher in Erken compared to Bolmen and Skogaryd) and nitrate (NO_3_-N) concentrations (higher in Bolmen and Skogaryd than in Erken). Moreover, the largest seasonal differences between summer and spring experiments in Bolmen and Erken were related to temperature, total phosphorus (TP), and chlorophyll *a* (chl-*a*) concentrations that were all lower in spring compared to summer. In contrast, Bolmen showed higher DOC, nitrate (NO_3_-N) and ammonium (NH_4_-N) concentrations in spring compared to summer (Table S1).

### Experimental additions of cDOM and inorganic N, P

As a standardized carbon source and browning agent, we used a peat extract solution prepared from commercially available unfertilized peat soil from a gardening centre (Langenheder et al. 2024). Peat soil was chosen to represent a natural cDOM source commonly found in northern temperate and boreal catchments in Fennoscandia, which typically contain organic-rich soils and peatlands (Pakarinen 1995). Similar cDOM extracts from commercially available peat soils have been used in mesocosm experiments before to successfully manipulate inputs of bioavailable allochthonous DOC and changes in light conditions (Gall et al. 2017, Mustaffa et al. 2020).

Separate extractions were done for the experiments in 2022 and 2023. Briefly, soil was homogenized and cold water extracted at elevated pH by adding 1M NaOH to maximize the extraction of dissolved organic matter (DOM) and other ions into solution phase. The pH was re-adjusted to neutral conditions using 1M HCl, the extract sequentially filtered through 0.2 μm, autoclaved, and aliquots of different volumes stored at -20°C. We added peat soil cDOM at a final concentration of 2 mg DOC L^-1^ to each mesocosm, which marks approximately the interval of differences between the 5^th^ and 95^th^ percentiles of DOC concentrations in Erken, and was chosen to present an increase that could easily occur with a single extreme event.

Inorganic P and N were added as K_2_HPO_4_ and NaNO_3_. The concentrations of the additions were based on data from a 30-year long time series from Erken, the only lake with data spanning several decades. For inorganic P, we added 45 μg PO_4_-P L^-1^, which corresponds to the 95^th^ percentile of all PO_4_-P concentrations measured in the lake since 1991. For the N additions, NO_3_ additions were chosen, as this is the main inorganic N compound that gets flushed into lakes via runoff and is the dominating form of N in inflowing water during peak flow periods (Hubbard et al. 2011, Miller et al. 2016). We added a standardized concentration of 400 μg NO_3_-N L^-1^, in both years (2022 and 2023), which corresponds to the maximum concentration measured historically in Erken. The PO_4_-P and NO_3_-N additions therefore represent conditions that could occur during a single extreme event, but still fall within the natural variability of concentrations found in the historical records (Langenheder et al. 2024). Together with smaller amounts of dissolved organic P and N from the peat extraction additions, this resulted in a total P and N addition of approximately 50 μg P L^-1^ (both years) and 500 and 460 μg N L^-1^ in 2022 and 2023, to each treated mesocosm, corresponding to a N:P supply ratio of approx. 22:1 and 20:1, respectively.

Sterile stock solutions of K_2_HPO_4_, NaNO_3_ and peat extract were prepared in batches that contained the exact mass by volumes for each addition to the mesocosms, which were then transported frozen to the different stations and thawed prior to each addition. More details can be found in Protocols 2 and 3 in Langenheder et al. (2024).

### High frequency monitoring

In each mesocosm, water temperature and dissolved oxygen concentrations were measured with an Oxygen Optode 4531 (Aanderaa Data Instruments AS, Norway), while chlorophyll-*a* (chl-*a*) and phycocyanin fluorescence and turbidity were measured by a Trilux sensor (Chelsea Technologies Group, UK). Both sensors were deployed at a 0.6 m depth for the duration of the experiment (Urrutia-Cordero et al. 2021b). Photosynthetically Active Radiation (PAR) expressed as photosynthetic Photon Flux Density (PFFD) was recorded at 0.4 m depth with an Apogee SQ-500 sensor (Urrutia-Cordero et al. 2021b). The PAR, Trilux and Oxygen/Temperature sensors were cleaned manually with a soft paint brush on a daily basis to avoid biofouling. All environmental parameters were measured with the lowest temporal resolution of 1 minute in Erken and Bolmen and 2 minutes in Skogaryd. All sensor data were processed in R version 4.3.2 (R Core Team 2023). Dissolved oxygen concentrations were calculated from measurements of oxygen saturation following Garcia and Gordon (1992). Quality control of sensor data was performed as follows: negative values were considered as erroneous sensor readings and removed from all data sets. Further outliers that noticeably exceeded peak concentrations were identified by visual inspection of the raw data and corresponding filtering thresholds for each environmental parameter in each year were specified and applied. Furthermore, consecutive repeats of identical measurements in the Trilux data were removed as erroneous sensor readings. Daily chl-*a* and phycocyanin concentrations were calculated from averaged fluorescence measurements taken between 23:00 and 03:00, which falls into the period between sunset (22:00) and sunrise (03:30) at the shortest day of the year at the northernmost site (Erken).

### Manual mesocosm sampling and measurements

The first sampling of the mesocosms was carried out on the day after they were filled, and then every fourth day until the end of the experiment. Prior to sampling, the mesocosms were gently mixed with a 20 cm diameter disc attached to a long wooden rod. Water samples were taken immediately after mixing from the middle of the enclosures at a depth of 0.6 m with a Ruttner sampler and filled into 2 L containers. Further sub-sampling for nutrient, chl-*a* analyses and absorbance measurements was conducted as described below. Additionally, a handheld multiprobe system (EXO2, YSI) was used in the Bolmen and Erken experiments and a Hach HQ2220 (Hach Lange AB, Helsingborg) multiprobe in the Skogaryd experiment to measure daily changes in pH and electrical conductivity and temperature at 0.6 m depth. In all experiments, PAR was recorded daily at three depths (0.05, 0.4 and 0.8 m) with a sensor with handheld display to determine the extinction coefficient. All sampling devices and handheld sensors were rinsed in the lake between sampling of different mesocosms.

Concentrations of DOC, TP, dissolved inorganic N (NO_3_-N, NO_2_-N, NH_4_-N), PO_4_-P and extracted chl-*a* in the water samples were measured at day 1 (before the first cDOM and nutrient additions), and then at days 5, 9, 13, 21 and 37 following standard protocols (Langenheder et al. 2024). Water colour (absorbance at 420 nm, a proxy for cDOM concentrations) was measured spectrophotometrically every fourth day. Lake water samples were taken outside the platform at a depth of 0.5 m.

### Calculation of metabolic rates

Metabolic rates (GPP, ER, and NEP) were calculated from the high-frequency dissolved oxygen measurements using the “LakeMetabolizer” R package (Winslow et al. 2016). Gaps in the sensor data were linearly interpolated if smaller than 40 minutes, and no metabolic rates were calculated on days with larger gaps. In addition to the sensor data, wind speed measurements were used to estimate surface gas exchange rates; in the case of Erken and Skogaryd, the wind speed was measured locally, and for Bolmen, these data were obtained from a meteorological station 17 km from the mesocosm facility. Wind speeds were measured at hourly or higher frequencies, but were linearly interpolated to match the sensor data.

The gas exchange velocity *k600* was calculated from wind speed using the method by Crusius & Wanninkhof (2003). As the mesocosms provided partial shelter from the wind, but were still affected by wave action, wind speeds were reduced by 50% before they were used in the metabolism calculations. Other percentages were tested as well, and although absolute metabolism estimates were affected, in most cases this did not strongly affect the treatment differences (Figure S1-S3). Thus, whilst the absolute values that we selected cannot be compared to other studies, the comparisons among mesocosms are robust.

The oxygen saturation concentration was calculated from water temperature based on Garcia & Gordon (1992), the “garcia-benson” model in LakeMetabolizer. A maximum likelihood metabolism model was then fitted based on *k600*, oxygen saturation, the depth of the mesocosm, and the observed oxygen concentrations, water temperature, and PAR (Winslow et al. 2016). The LakeMetabolizer package provides unconstrained estimates of metabolic rates, meaning that physically impossible (negative) values of GPP and ER can be generated, due to noise in the data, sensor errors, or a dominating physical signal compared to a biological signal. We decided to retain rates between 0 and -0.5 mg O_2_ L^-1^ d^-1^ (where a value of 0.5 mg O_2_ L^-1^ d^-1^ represents ca. 50% of and 60% of the average GPP and ER for all lakes, respectively) as these rates could be due to noise close to a mean of 0, which is possible at low biological levels. In contrast, we removed values below -0.5, as these were more likely due to invalid measurements. In that way, on average 1.6% of estimates of GPP and 2.2% of ER were removed from the entire dataset.

### Statistical analysis

The responses of different variables to the experimental treatments (Control, Daily, Intermittent, Extreme) were tested by Generalized Additive Mixed Models (GAMM; Wood 2010) using the "mgcv" package (Wood 2011) in R version 4.3.2 (R Core Team 2023) which enables capturing curvilinear responses (Zuur et al. 2009, Carvalho et al. 2013). Different models were tested and compared using AIC and BIC to select the most parsimonious model. For sensor measurements, average daily values were calculated prior to the statistical analysis.

The final model included an interaction term between Experiment, Day and Treatment with a smoothing factor (s), Mesocosm (1-16) nested in Experiment (Bolmen summer, Bolmen spring, Erken summer, Erken spring and Skogaryd) as random effects, and Experiment Day in a corAr1 argument to account for the autocorrelation structure in time series data. The global data from the automated sensors (chl-*a* and phycocyanin fluorescence, dissolved oxygen concentration and PAR) were too noisy to fit GAMMs with multiple random factors, so Experiment and mesocosm were combined into one random factor (ID). All analyses were performed based on a normal (Gaussian) distribution and identity link function, which resulted in satisfactory normal distribution of the residuals for all models. The GAMMs were run for global datasets including all lakes and for data from individual experiments (to test whether they followed the same trend as the overarching models or exhibited different responses). We also ran separate analyses for all individual experiments in cases when there were no significant differences between treatments in the global model to investigate if single lake experiments showed opposing patterns that cancelled each other out. Estimated marginal means and pairwise contrasts among the individual interaction terms (treatments over time) were computed for the entire experiment and for individual time points using the R package "emmeans" (Lenth 2024). The *p*-values were adjusted for multiple comparisons using the Tukey method. Pairwise comparisons were visualized with heat maps.

## Results

The GAMM models showed statistically significant effects of most smoothed terms on all different response variables, i.e. most environmental conditions that we expected to be directly affected by the treatment manipulation (water colour, NO_3_-N, PO_4_-P, TP and DOC), chl-*a* and phycocyanin fluorescence, GPP, ER and NEP (Table S2). This included significant effects of the Treatment-Day interaction as well as random effects of mesocosm ID and experiment (in case of the global model based on all experiments together) (Table S2).

### Treatment effects on environmental conditions

There was clear temporal variation in water colour, DOC, NO_3_-N, and TP concentrations in all treatments during the experiments, while PO_4_-P concentrations increased only in the Extreme treatment (Figs S4-S8). DOC and PO_4_-P concentrations also changed in the Control over time but this was less pronounced compared to the runoff treatments and pairwise comparisons in the overarching model revealed significantly higher water colour, TP, and NO_3_-N concentrations over the course of the experiments in all runoff treatments compared to the control (Table S3). Further, DOC concentrations in the Extreme and Intermittent treatments and PO_4_-P in the Extreme treatment were significantly higher compared to the Control in the overarching models and in two of the experiment-specific models (Erken spring and Skogaryd). There were also differences in environmental conditions among the runoff treatments. Overall, the Extreme treatment had higher PO_4_-P and NO_3_-N concentrations than the Daily and Intermittent treatments reflecting the steep increase in concentration after the addition on Day 8 (Figs S6 and S8). The experiment specific models gave similar results for NO_3_-N in all cases, even though the increase between the Extreme and Intermittent treatment was not statistically significant in the Bolmen summer experiment. For PO_4_-P, increases were only statistically significant in the global model, in Skogaryd, and for Erken in spring. TP concentrations were higher in the Extreme and Intermittent treatment compared to the Daily treatment, even though this was only significant in case of the overarching model and the Bolmen summer experiment. In the Bolmen spring experiment TP concentrations were also higher in the Extreme than in the Intermittent treatment. Finally, TP concentrations were in all cases higher in the runoff treatments than in the Control (Table S3, Fig S7).

At the end of the simulated runoff period (Day 20), PO_4_-P and NO_3_-N concentrations converged across all runoff treatments and subsequently reached similar or lower concentrations than the Control by the end of the experiment (Day 37) (Figs S6 and S8). On the contrary, water colour also converged among the runoff treatments until the end of the simulated runoff period, but remained higher than in the Control throughout the recovery period in all runoff treatments (Fig S4). In all experiments, except Erken spring, TP concentrations remained higher in the runoff treatments compared to the control, whereas differences between the runoff treatments were small at the end (Fig. S7).

Additional information on treatment effects on other measured environmental variables (PAR, NO_2_-N, NH_4_-N, chl-*a* concentration from extracted filters, conductivity and pH) can be found in the Supplementary material (Text S1).

### Changes in chl-*a* and phycocyanin concentrations

Across all lakes and seasons, chl-*a* and phycocyanin concentrations changed significantly over time in the Daily, Intermittent and Extreme treatment, but not in the Control (Figs 2 and 3, Table S2). For the overarching model using the combined dataset, pairwise comparisons showed that chl-*a* concentrations in the Extreme treatment were higher compared to all other treatments over the course of the experiment (Table 1). More specifically, chl-*a* concentrations peaked in the Extreme treatment towards the end of the simulated runoff period when they became higher compared to the Daily and Intermittent treatments (Fig. S10). The same pattern was observed in the experiment specific analyses in the Erken and Skogaryd experiments (Fig. 2, Table 1), though differences in the latter were not significant compared to the Intermittent treatment and only marginally significant compared to the Daily treatment (Fig. 2). Chl-*a* concentrations in the Intermittent treatment were higher compared to the Control in the overarching model and in the specific model for Skogaryd, lower compared to the Control in Erken summer and higher compared to the Extreme and Daily treatment in Bolmen summer. Chl-*a* concentrations in the Daily treatment did not differ from the Control, except in Skogaryd. A closer inspection of the data and model results still revealed that chl-*a* concentrations increased significantly in the Daily and Intermittent treatment in the overarching model, as well as in the two Erken, summer Bolmen and Skogaryd experiments during or after the simulated runoff period, and later decreased to or below levels in the Control. Skogaryd remained different compared to the Control for the longest period of time (Fig 2, Fig. S10).

**Figure 2.**
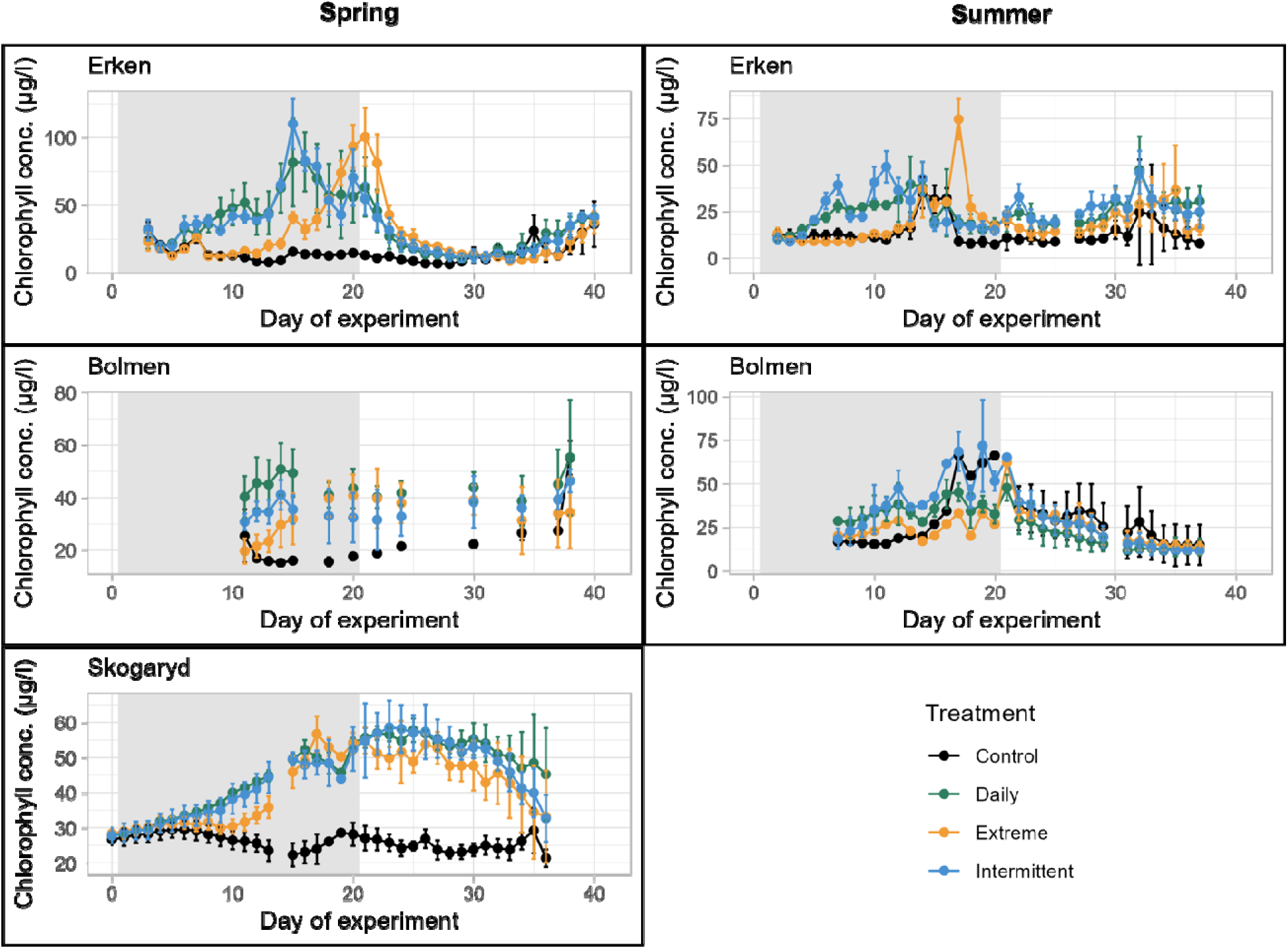
Changes in chl*-a* concentration over time calculated from daily average of high-frequency chlorophyll *a* fluorescence measurements in the mesocosm experiments done in spring 2023 and summer 2022 in Erken, Bolmen and Skogaryd. The shaded area in grey is the simulated runoff period in which nutrients were added (see Figure 1). Note the differences in the y-axis scale limits between graphs.

**Figure 3.**
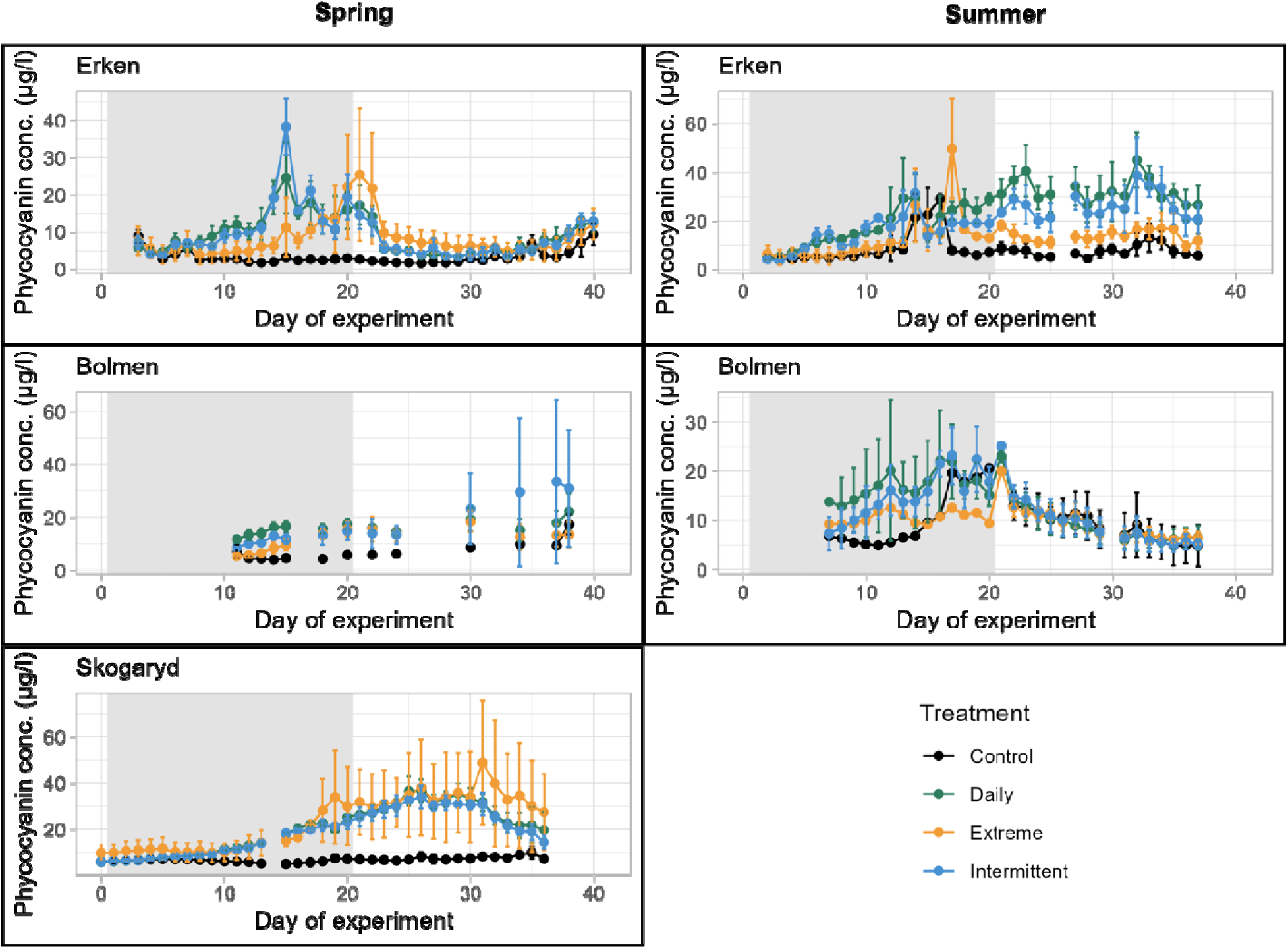
Changes in phycocyanin concentration over time, calculated from daily average of high-frequency phycocyanin fluorescence measurements in the mesocosm experiments done in spring 2023 and summer 2022 in Erken, Bolmen and Skogaryd. The shaded area in grey is the simulated runoff period in which nutrients were added (see Figure 1). Note the differences in the y-axis scale limits between graphs.

**Table 1.**
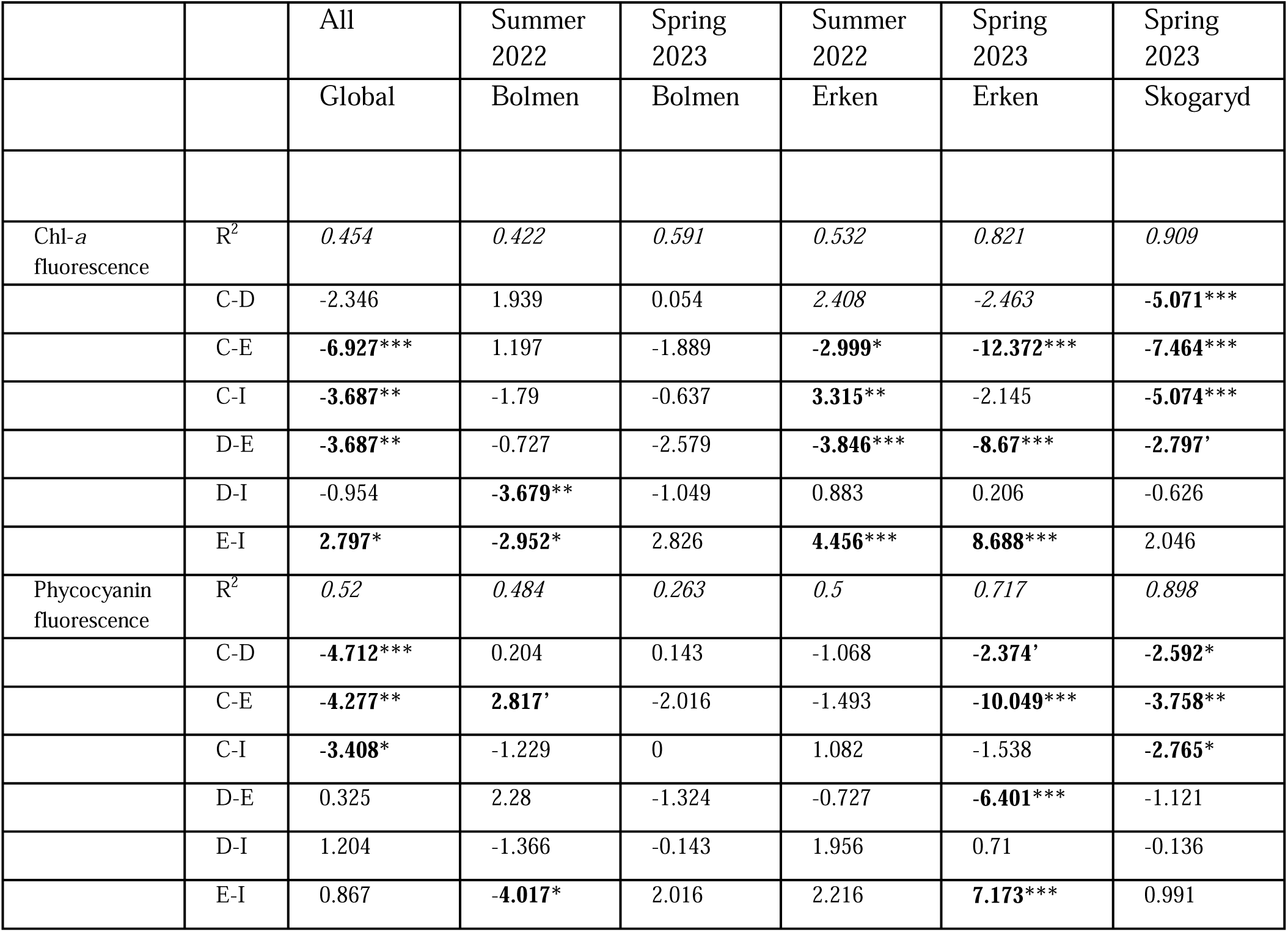
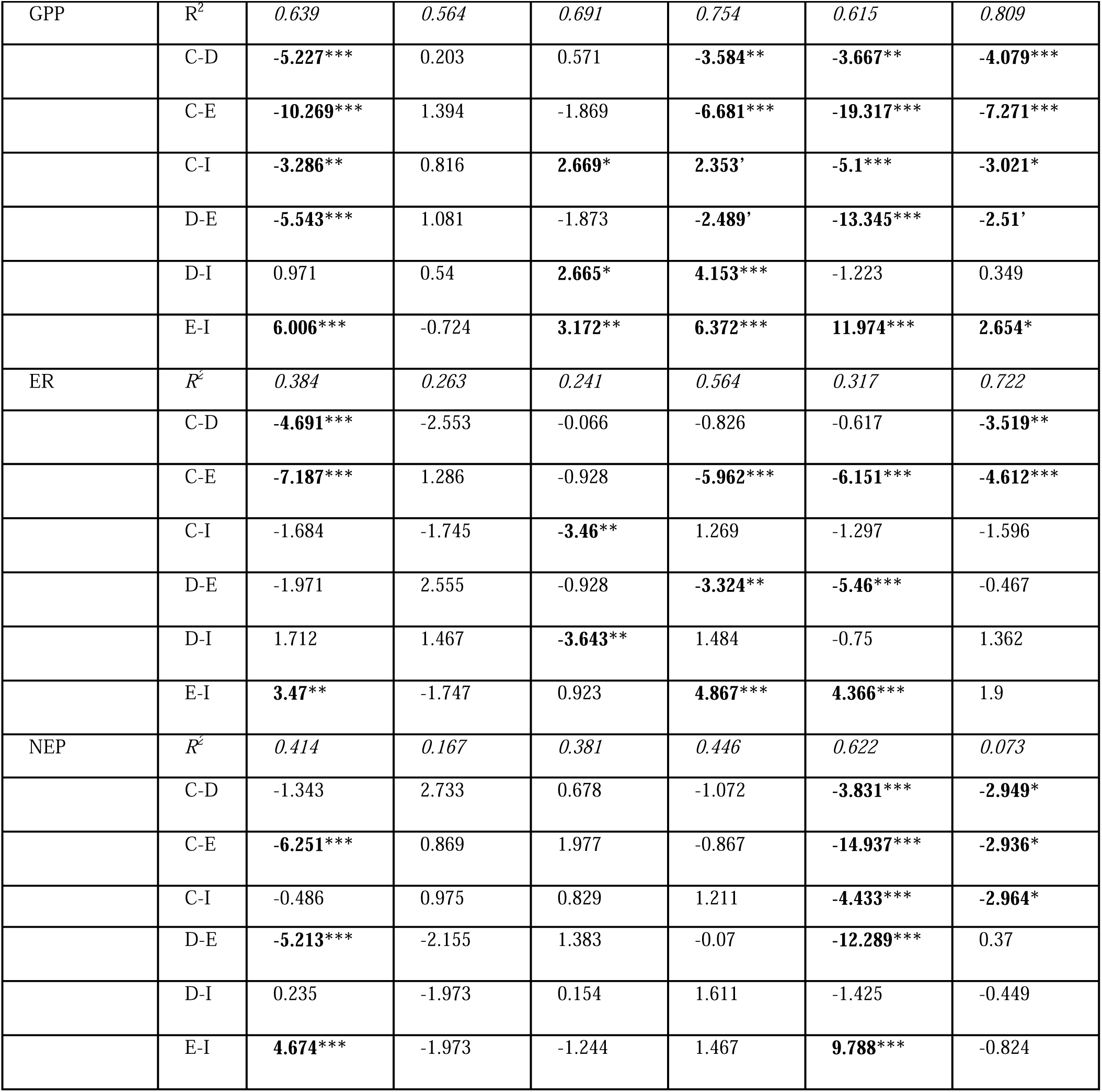
R^2^ values of single lake models and tT-ratios of pairwise comparisons of interaction terms of treatments and time in GAMMs based on all experiments or separate single experiments. C: Control, D: Daily Treatment, E: Extreme treatment, I: Intermittent treatment. Response variables tested were daily averages of chlorophyll *a* and phycocyanin concentrations (μg L^-1^) from fluorescence measurements, and GPP, R, and NEP (μg O_2_ L^-1^ day^-1^) calculated from day and nighttime changes in dissolved oxygen concentrations. R-sq: adjusted R^2^ values, i.e. the proportion of variance explained by the model. Significance codes: *** < 0.001, ** <0.01, * < 0.05, ‘ < 0.1. All values that follow the significance terms above are highlighted in bold. F-statistics and degrees of freedom of the models can be found in Table S3.

Across all lakes and seasons, phycocyanin concentrations were higher in the runoff treatments than in the Control treatment and showed similar temporal dynamics than chl-*a* concentrations across treatments (Table 1, Fig. 3, Fig. S11). Most remarkably, the Erken spring experiment showed the same ‘delayed peak’ in phycocyanin concentrations as measured for chl-*a* concentrations in the Extreme treatment, and consistently higher concentrations compared to all other treatments (Fig. 3, Fig. S11, Table 1). Phycocyanin concentrations in the Extreme treatment were higher compared to the Intermittent treatment in all cases (albeit only statistically significantly in Erken spring), but lower in Bolmen spring.

By the end of the experiment (day 37) there were in most cases no differences in chl-*a* and phycocyanin concentrations between the runoff treatments and the Control and among runoff treatments (Figs 2 and 3, Figs S10 and S11).

### Changes in metabolic rates

GPP and ER changed in all treatments including the Control throughout the experiment (Table S2). The same was the case for NEP with the exception of the Daily treatment.

GPP was consistently higher in the Extreme than in all other treatments when all experiments were analysed together (Table 1). Moreover, GPP was higher in the Intermittent and Daily treatments compared to the Control. Both Erken and the Skogaryd experiments followed this general trend, albeit differences between runoff treatments were less pronounced in the case of the latter (Fig. 4). In the Erken experiment, GPP switched from being lower to higher in the Extreme compared to the other runoff treatments approximately 1 week after the pulse addition, an effect that lasted for 6 days (Fig. 4, Fig. S12). In the Bolmen spring experiment, GPP in the Intermittent treatment was lower compared to all other treatments during most of the simulated rainfall period, but higher during the last week of the experiment (Fig. 4, Fig. S12). In the Bolmen summer experiment, there were no differences in GPP between treatments (Table 1).

**Figure 4.**
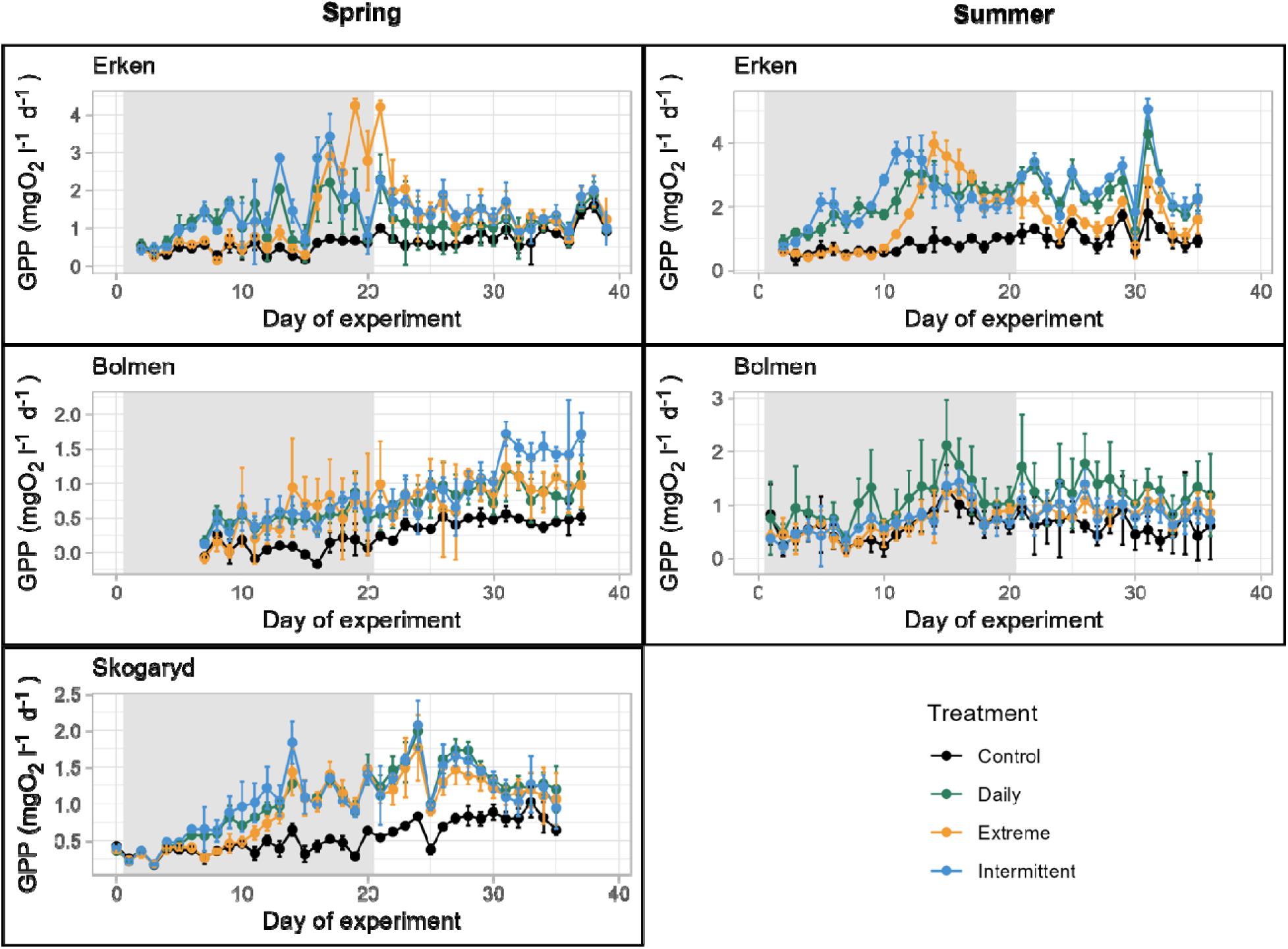
Changes in GPP over time, calculated from daytime changes of dissolved oxygen concentration from high-frequency sensor measurements in the mesocosm experiments done in spring 2023 and summer 2022 in Erken, Bolmen and Skogaryd. The shaded area in grey is the simulated runoff period in which nutrients were added (see Figure 1). Note the differences in the y-axis scale limits between graphs.

Across all experiments, ER was higher in the Extreme than in all other treatments, albeit this difference was not statistically significant when compared to the Daily treatment (Table 1). Further, ER was higher in the Daily treatment than in the Control. The overall trend that ER was higher in the Extreme treatment compared to all other treatments was seen in both Erken experiments, but was less clear in the other experiments (Table 1, Fig. 5). Likewise, NEP was also significantly higher in the Extreme treatment compared to all other treatments across all experiments and this trend was strongly driven by the Erken spring experiment (Table 1, Fig. 6). Similar NEP and ER also switched most pronouncedly in the two Erken experiments from being lower in the Extreme compared to the Intermittent and Daily treatments in the beginning, to being higher at later stages of the simulated rainfall period (Figs 5 and 6, Figs S13 and S14).

**Figure 5.**
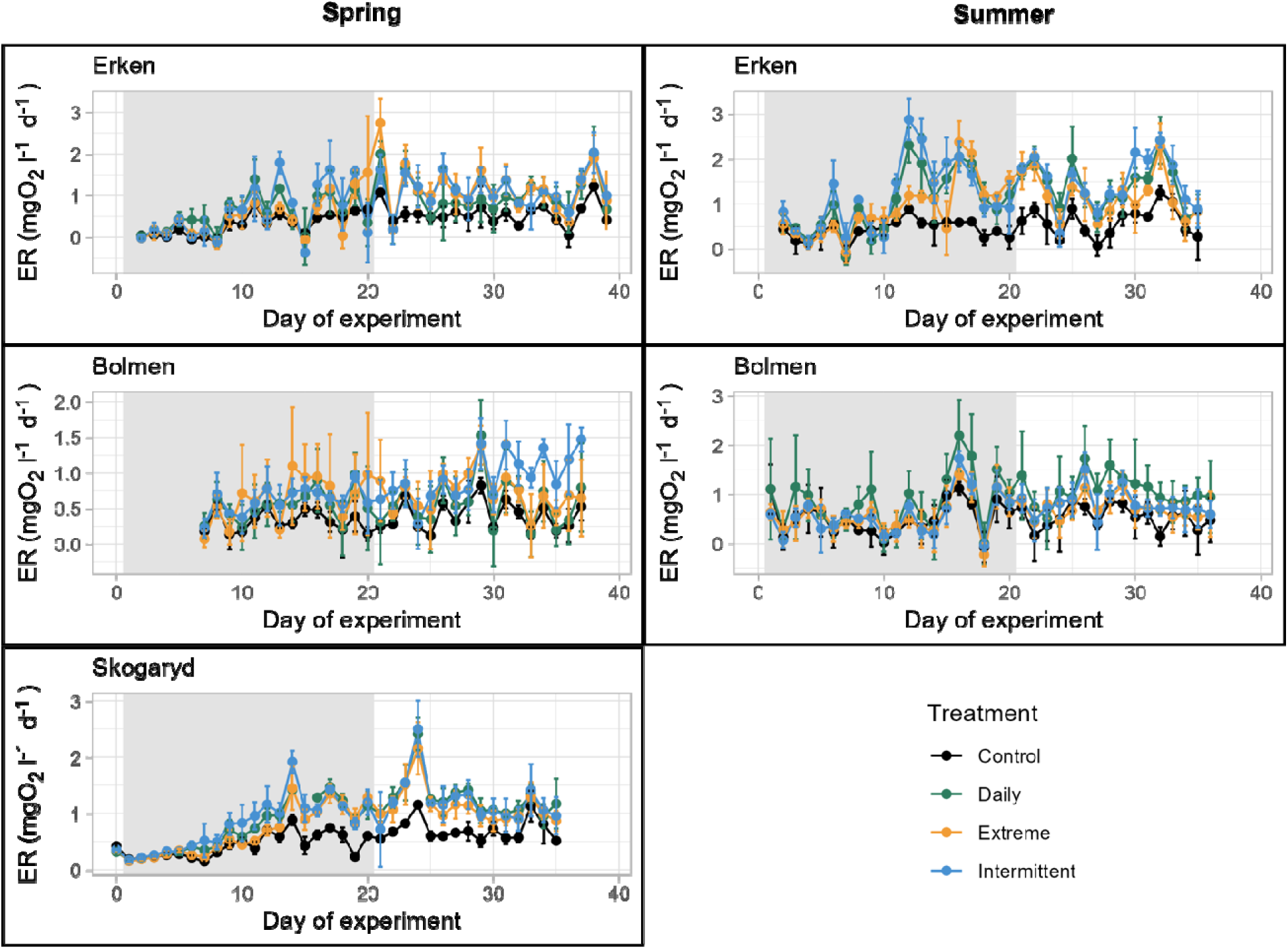
Changes in ER over time, calculated from daytime changes of dissolved oxygen concentration from high-frequency sensor measurements in the mesocosm experiments done in spring 2023 and summer 2022 in Erken, Bolmen and Skogaryd. The shaded area in grey is the simulated runoff period in which nutrients were added (see Figure 1). Note the differences in the y-axis scale limits between graphs.

**Figure 6.**
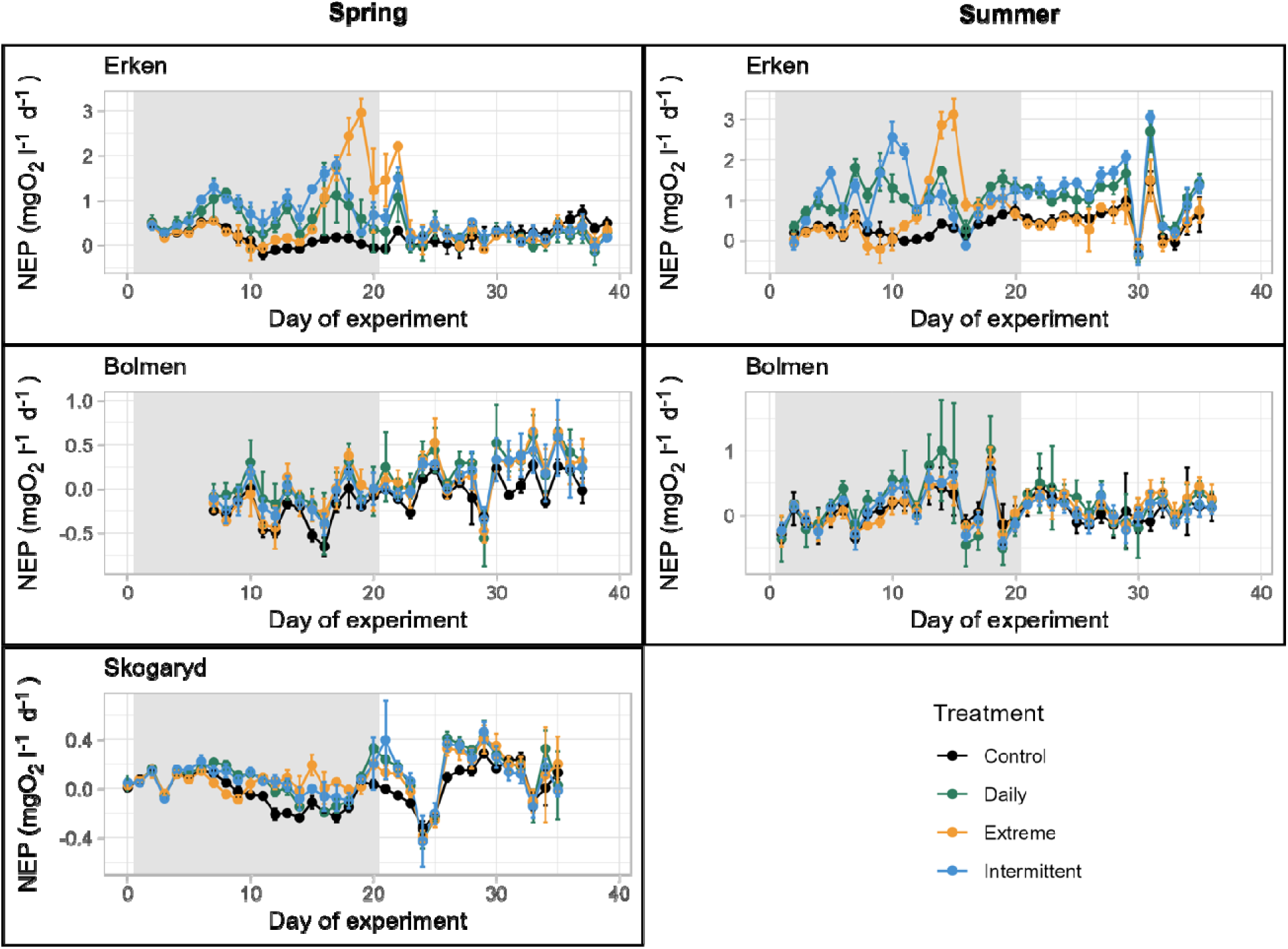
Changes in NEP over time in the mesocosm experiments done in spring 2023 and summer 2022 in Erken, Bolmen and Skogaryd. The shaded area in grey is the simulated runoff period in which nutrients were added (see Fig. 1). Note the differences in the y-axis scale limits between graphs.

By the end of the experiment (day 37), all metabolic rates in the runoff treatments were similar or even lower than in the Control in most cases (Figs S10-14). There were also only few significant differences between treatments except GPP and ER being higher in the Intermittent compared to the Daily and Extreme treatments in the Bolmen spring experiment (but note that the R^2^ values of GAMMs were low in both cases, Table 1, Figs S12-14).

## Discussion

We conducted a mesocosm experiment, replicated across space (three lakes) and time (two seasons in two of the lakes), to compare different runoff scenarios in which plankton communities received the same total amount of cDOM and nutrients, but at different pulse intensity and frequency. Overall, we found that differences in runoff pulse characteristics affected chl-*a* and phycocyanin concentrations and metabolic rates differently, and that more pronounced effects occur in response to the extreme pulse scenario compared to scenarios with more frequent, smaller pulse additions. Moreover, the effects differed between lakes, but less so between seasons, and were in most cases transient.

The different runoff treatments led to overall consistent changes in dissolved P and N concentrations across lakes and seasons, reflecting differences in the magnitude and frequency of the applied cDOM and inorganic nutrient additions, with the overall largest differences in the Extreme treatment where cDOM and nutrients were added in one large pulse. While NO_3_-N concentrations remained higher compared to the control in the runoff treatments after the simulated runoff period, especially in those lakes with higher background N concentrations (Bolmen and Skogaryd), PO_4_-P concentrations dropped rapidly to background levels in all experiments. This suggests overall strong P limitation, which was expected as the background N:P ratios calculated using dissolved total N and TP concentrations were > 50:1 in all cases. In addition, the results also point to more rapid biological uptake of PO_4_-P than NO_3_-N in Bolmen and Skogaryd, while in Erken both PO_4_-P and NO_3_-N were rapidly utilized, suggesting a stronger initial N limitation in the case of Erken. As expected, water colour increased in the runoff treatment compared to the Control in congruence with the pulse addition scenarios and the same was found for DOC concentrations except in the summer experiment in Erken. This exception could point to higher and more rapid degradation of the added allochthonous DOC through bacterial degradation (Tranvik 1992) and photodegradation (Dempsey et al. 2020), or both. In congruence with the latter idea, PAR reached the highest values in the Erken summer experiment (Fig S9), and therefore possibly also UV radiation, which could have promoted photodegradation and increased bioavailability of the added allochthonous DOC (Bertilsson and Tranvik 1998, Cory and Kling 2018, Dempsey et al. 2020).

We found that adding the same amount of cDOM, N and P at pulses of different magnitude, number and frequency caused differences in both chl-*a* and phycocyanin concentrations and metabolic rates (GPP, ER, NEP), with the overall strongest increase and deviating temporal dynamics when cDOM and nutrient additions occurred as one large pulse compared to the scenarios with smaller, more regular additions. There was a more gradual increase in phytoplankton parameters in the Daily and Intermittent treatments in the simulated rainfall period, whereas the Extreme treatment had a delayed effect, which happened several days after the large pulse addition resulting in more rapid growth. This pattern was very noticeable in the clearwater lake (Erken), but less so in the other two lakes, particularly in Bolmen, demonstrating that ecosystem responses to pulse additions were context dependent. At the end of the recovery period, almost all runoff treatments converged to the levels of the Control. Similar transient effects and recovery from large nutrient pulses and extreme precipitation events have also been found in other studies (Cottingham and Schindler 2000, de Eyto et al. 2016). Overall, phycocyanin concentration increased in the runoff treatments in all experiments, which supports previous studies showing that cyanobacteria can be favoured by increases in N and P concentrations as well as decreases in light availability associated with higher cDOM concentrations (Creed et al. 2018, Reinl et al. 2021). There are, however, also examples in the literature where browning suppressed cyanobacteria despite high nutrient availability (Lyche Solheim et al. 2024) and it has also been found that low levels of browning promote cyanobacterial growth and higher levels that of cryptophytes (Senar et al. 2021). Further studies are therefore needed to investigate interactions between browning-induced differences in light levels and other factors (e.g. nutrient concentration and stoichiometry) on cyanobacterial growth.

In general, pulse additions led to stronger increases in GPP than in ER, resulting in net autotrophic systems (i.e. more positive NEP). This finding suggests that stimulating effects of inorganic nutrient additions outweighed potential suppressing effects related to light limitation and phytotoxic effects caused by increased browning from the cDOM addition (Gjessing and Kallovist 1991, Rodríguez et al. 2016, Zwart et al. 2016, Kelly et al. 2018). As reported in other lake studies, changes in ER were coupled to those of GPP, so that increases in GPP were accompanied by increases in ER (e.g. Solomon et al. 2013, Scharfenberger et al. 2019). In our case, this was most clearly seen in single lake experiments where the strongest changes in ER were found when also GPP also changed most strongly. This pattern suggests that ER increased through phytoplankton-derived autochthonous DOC as well as increasing phytoplankton respiration, in combination with bacterial degradation from the added DOC source (Attermeyer et al. 2014, Pace et al. 2021, Calderó-Pascual et al. 2022).

The applied cDOM and nutrient additions led to overall increases in NEP in all the study lakes regardless of their background conditions. This could be partially explained by the relative increases in concentrations, which were stronger for nutrients compared to DOC. Different supply ratios between DOC and inorganic nutrients can also lead to different effects on GPP and thereby NEP (Kelly et al. 2018, Olson et al. 2020) and increases in ER have also been found in response to cDOM inputs and extreme precipitation events (Klug et al. 2012, Sadro and Melack 2012, de Eyto et al. 2016). In addition, even though previous experiments using the same extraction procedure from commercially available peat soil indicated that the extracted DOC is bioavailable (Gall et al. 2017), we lack specific information about its bioavailability in our experiment, making it difficult to evaluate its potential to enhance bacterial growth and ecosystem respiration. Hence, similar experiments to ours should be carried out in the future with different DOC concentrations or a different type of DOC source, e.g. more bioavailable leaf litter extract (Brett et al. 2017). Despite constraints related to the type of cDOM addition, the main conclusion remains valid: it is not only the overall magnitude of cDOM and inorganic nutrient inputs, but also their frequency and timing, that need to be considered when predicting ecosystem impacts of runoff events.

As hypothesized, we found that the effects of the runoff treatment on chl-*a* and phycocyanin concentrations and metabolic rates differed between lakes. In contrast, seasonal differences within lakes were less pronounced. This suggests that background chemical characteristics, such as water colour, have a stronger effect on phytoplankton and cyanobacterial biomass and metabolic rates than seasonal differences in e.g. plankton composition and nutrient concentrations within lakes. There were clear differences between the Extreme and the other two runoff treatments in Erken, but not in Bolmen and Skogaryd, demonstrating that ecosystem responses to different intensities and frequencies of pulse additions were modulated by local conditions. Similarly, Thayne et al. (2022) found that variation and resistance to storm impacts depended more on antecedent conditions in the lake prior to the storm, including turbidity, light conditions and water chemistry, than on actual storm characteristics. This is consistent with the differing responses of chl-*a* and phycocyanin concentrations and metabolic rates among the lakes, which likely arose from contrasts in light limitation and nutrient availability driven by their background cDOM and NO_3_-N concentrations. In addition, varying sensitivities of the lake communities to the applied cDOM and nutrient additions might also have played a role. Hence, it could be possible that an overall lower light limitation in Erken and potentially stronger initial N limitation, might have resulted in a more adaptive phytoplankton community in response to the extreme pulse. Another possible explanation could be that bacterial communities adapted to more humic waters in Bolmen and Skogaryd outcompeted phytoplankton, because they were more strongly stimulated by peat extract additions. If the latter mechanism was important, we would, however, have expected that enhanced bacterial production should at least have led to a stronger increase in respiration in the Extreme treatments in Bolmen and Skogaryd, which was not the case. Less pronounced treatment responses in Bolmen and Skogaryd compared to Erken could, nevertheless be due to the fact that additions were chosen based on historical records from Erken and designed in a way that the Extreme treatment would represent historically rare nutrient inputs as well as significant variability in DOC concentration compared to average concentrations. For the other lakes, however, these assumptions might not hold and communities in those lakes might have been more resistant to the applied changes.

In addition, Erken showed stronger relative and absolute changes for several response variables between the runoff treatments and the control compared to the humic lakes (Bolmen and Skogaryd). This was expected given that primary production typically becomes increasingly constrained by light limitation along gradients of increasing water colour and cDOM concentrations (Thrane et al. 2014, Seekell et al. 2015, Deininger et al. 2017, Isles et al. 2021). At the same time, clear and significant differences in chl-*a*, phycocyanin concentrations, and GPP were found between the runoff treatments and the Control in the Skogaryd experiment, whereas this was not found in the two Bolmen experiments. This was surprising, because we expected the high humic content in the lake at Skogaryd and the resulting light limitation to constrain any stimulating effects of the nutrient additions. One possible reason could be that phytoplankton species in Skogaryd are well adapted to low-light conditions and therefore positively respond to pulses of cDOM and nutrients. It also suggests that other factors than light limitation constrained phytoplankton responses in the Bolmen experiments, such as stronger growth limitation by other nutrients than N and P (e.g. silica), differences in zooplankton grazing and levels of adaptation of phytoplankton communities to environmental variability. Moreover, periphyton biomass on polyethylene strips that were employed in the mesocosms during the experiments was higher in the runoff experiments compared to the Control in Bolmen, albeit this was only significant in the spring experiment (Text S2, Fig. S18). This indicates that a larger amount of dissolved nutrients was channelled into benthic biomass production, constraining pelagic growth more strongly compared to the Skogaryd and Erken experiments. Interestingly, there were significant effects on chl-*a* concentrations measured on a few selected time points by spectrophotometric pigment analyses, whereas differences in chl*-a* concentrations based on fluorescence measurements were not significant. It could be that the numerous gaps in the sensor data provide a less reliable estimate of chl-*a* concentrations based on chl-*a* fluorescence.

However, given that weak and non-significant effects were also found for other response variables irrespective of the number of underlying datapoints, the conclusion that Bolmen showed the weakest overall response to the runoff treatments out of the three lakes remains valid.

Chl-*a* concentrations from sensor measurements were in several cases higher than those obtained from extracted samples. Chl-*a* fluorescence is influenced by, among others, phytoplankton composition as well as cDOM concentrations (Lawrenz and Richardson 2011, Kuha et al. 2020) and can overestimate chl-*a* concentrations. It is therefore recommended to calibrate chl-*a* fluorescence signals with measurements of extracted concentrations (Levi et al. 2025). We did not do this here due to the low number of samples in case of the latter, but also because it was recently found that both chl-*a* estimates can show different correlations with phytoplankton biomass measured by microscopic counts, suggesting that they should be considered as complementary and not replaceable measurements of phytoplankton biomass (Urrutia-Cordero et al. 2024). Nevertheless, any biases related to chl-*a* fluorescence measurements should not affect the conclusions of our study, which was focused on the comparison of effects between treatments.

To conclude, our results suggest that extreme cDOM and nutrient pulses have the potential to disrupt lake ecosystem responses compared to scenarios with less extreme and more gradual inflows, which is important as extreme weather events are expected to become more intense and frequent in the future as a result of climate change (Jennings et al. 2012, Zwart et al. 2017, Stockwell et al. 2020). Importantly, these effects were transient: differences among runoff treatments as well as differences compared to the Control generally vanished in a few weeks.

Our study also showed that responses differ among lakes, highlighting the importance of context dependency. More specifically, the strongest effects between different runoff scenarios were found in the clearwater lake (Erken), whereas lakes with higher background water colour and NO_3_-N concentrations (Bolmen, Skogaryd) were less affected. This finding suggests that these two environmental variables, water colour and NO_3_-N concentrations, could be used as indicators of the vulnerability of boreal lake ecosystems to extreme runoff events. We therefore encourage future studies to use the standardized protocol that was established as part of this study, which allows straightforward replication of the experiment in additional lakes. This would create a larger data set that can be used to directly decipher and quantify the underlying drivers behind variable responses between lakes using machine learning tools and structural equation modelling (Lefcheck 2016, Su et al. 2024). It is important to keep in mind that we only applied one extreme pulse in our experiment and it has been shown that repeated extreme events can have long-term effects on phytoplankton biomass and metabolic rates (e.g. Stelzer et al. 2022).

Another venue for future experimental studies should also address the effects of repeated events and focus on how differences in stoichiometric nutrient ratios between catchment inputs and *in situ* ratios in lakes can also affect lake metabolism (Corman et al. 2023). This would be particularly valuable if, as also done here, such experiments are connected with historical data from long-term monitoring programmes so that the effect of realistic scenarios can be addressed. Finally, extreme weather events such as storms, can also influence physical lake properties that were not addressed in our study, including thermocline deepening, temperature changes and dilution effects, which can also influence phytoplankton biomass and metabolism (Giling et al. 2017, Doubek et al. 2021, Grossart et al. 2025). Studies that lead to a more integrative understanding of multiple effects of runoff loading on plankton communities are therefore needed as well.

## Acknowledgements

This project was supported by the EU H2020-INFRAIA project (871081) AQUACOSM-plus—Network of Leading Ecosystem Scale Experimental AQUAtic MesoCOSM Facilities Connecting Rivers, Lakes, Estuaries, and Oceans in Europe and beyond—funded by the European Commission, both for salary to PIs and technical staff which supported SAB, JCN, SL, IS, NK, WCM and through its Transnational Access program, which supported A.A., Su.B., Ben.B., E.A.C., N.Car., N.Cat., P.E.A., I.E., B.G., E.G., Nu.K., A.K., A.L., C. M-L., A. P-M., N.P., M.M.Y., and K.Y. This study was further supported by the Swedish Research Council for the Swedish Infrastructure for Ecosystem Science (SITES), in this case at the Erken Laboratory and the Skogaryd Research Catchment (Swedish Research Council grant 2021-00164). J.P.M. was funded by Formas (Swedish Research Council for Sustainable Development) and the European Union’s Horizon Europe Programme under the 2022 Joint Transnational Call of the European Partnership Water4All Grant Agreement no. 101060874 (MEWS project) and Horizon 2020 Programme Grant Agreement no. 101017861 (SMARTLAGOON project).

M.S, C.M. and J.E were supported by the German Research Foundation (DFG, STR 1383/5-1 and STR 1383/8-1). A.H. was supported by a DAAD (German Academic Exchange Service) research grant for doctoral candidates (No. 57556281). J.S. and K.J. was supported by the Swedish Research Council FORMAS (2020–00730) and B.B. was supported by the Swedish Research Council VR (2019–03970). I.E. was funded by FRESH - NERC Centre for Doctoral Training in Freshwater Biosciences and Sustainability [NE/R011524/1]. Su.B. was supported by the projects EVASIONA (PID2021-122817NB-100) and RIPAMED (CNS2023-144737) funded by MICIU/AEI/10.13039/501100011033 and NextGenerationEU/PRTR.

## Data availability statement

All data will be made available in the SITES Dataportal and R scripts used for the calculations of metabolic rates on the SITES GitHub.

## Author contributions

**Silke Langenheder:** Conceptualization, Data Curation, Funding Acquisition, Investigation, Project Administration, Writing – original draft preparation, Writing – Review & Editing; **Jorrit Mesman**: Conceptualization, Formal Analysis, Visualization, Writing – original draft preparation, Writing – Review and editing; **Nils Kreuter**: Data Curation, Investigation, Project Administration, Writing – original draft preparation, Writing – Review & Editing; **Dolly Kothawala**: Conceptualization, Investigation, Project Administration, Writing – Review & Editing; **Gabriela Ágreda-López**: Investigation, Writing – Review & Editing; **Akif Ari**: Investigation, Writing – Review & Editing; **Stella A Berger**: Conceptualization, Funding Acquisition, Investigation, Project Administration, Writing – Review & Editing; **Susana Bernal**: Investigation, Writing – Review & Editing; **Bence Buttyán**: Investigation, Writing – Review & Editing; **Berenike Bick**: Investigation, Writing – Review & Editing; **Nuria Carabal**: Investigation, Writing – Review & Editing; **Nuria Catalan**: Investigation, Writing – Review & Editing; **Eleni A Charmpila**: Investigation, Writing – Review & Editing; **William Colom Montero**: Data Curation, Investigation, Writing – Review & Editing; **Pelin Ertürk Arı**: Investigation, Writing – Review & Editing; **Inge Elfferich**: Investigation, Writing – Review & Editing; **Johanna Exner**: Investigation, Writing – Review & Editing; **Bence Gergácz**: Investigation, Writing – Review & Editing; **Emma Gray**: Investigation, Writing – Review & Editing; **Anika Happe**: Conceptualization, Investigation, Writing – Review & Editing; **Congcong Jiao**: Investigation, Writing – Review & Editing; **Kevin Jones**: Investigation, Writing – Review & Editing; **Nusret Karakaya**: Investigation, Writing – Review & Editing; **Antonija Kulaš**: Investigation, Writing – Review & Editing; **Anna Lupon**: Investigation, Writing – Review & Editing; **Clara Mangold**: Investigation, Writing – Review & Editing; **Clara Mendoza-Lera**: Investigation, Writing – Review & Editing; **Jens Nejstgaard**: Conceptualization, Funding Acquisition, Investigation, Project Administration, Writing – Review & Editing; **Jimmy C. Oppong**: Investigation, Writing – Review & Editing; **Angela Pedregal-Montes**: Investigation, Writing – Review & Editing **Nuria Perujo**: Investigation, Writing – Review & Editing; **Juha Rankinen**: Project Administration, Writing – Review & Editing; **Tobias Rütting**: Project Administration, Writing – Review & Editing; **Johanna Sjöstedt**: Conceptualization, Investigation, Project Administration, Writing – Review & Editing; **Maren Striebel**: Conceptualization, Project Administration, Implementation, Writing – Review & Editing; **Katerina Symiakaki**: Investigation, Writing – Review & Editing; **Eline van Dam**: Investigation, Writing – Review & Editing; **Simon Wentritt**: Investigation, Project Administration, Writing – Review & Editing; **Majd M Yaqoob**: Investigation, Writing – Review & Editing; **Kadir Yıldız**: Investigation, Writing – Review & Editing, **Ingrid Sassenhagen**: Conceptualization, Data Curation, Investigation, Project Administration, Formal analysis, Writing – original draft preparation, Writing – Review and editing.

