## Supplementary files for "Lake ecosystem responses to runoff variability across time and space"

Supplementary text.

**Text S1. Treatment effects on $NO_2$-N, $NH_4$-N, and chl-*a* concentration from extracted filters, conductivity and pH.**

There were significant changes in all parameters in all treatments throughout the experiments, except $NH_4$-N concentrations in the Control and Daily treatments (Table S2). Overall $NO_2$-N and $NH_4$-N concentration were low in all experiments (Figs S15 and S16) and were close to detection limit in all cases. For pairwise comparisons the overarching GAMM models as well as the experiment specific models for Bolmen spring, Erken summer and Erken spring showed that *chl-a* concentrations in samples extracted from filters were higher in the Daily and Intermittent treatments compared to the Control during the course of the experiment (Fig. S17, Table S3). Further they were higher in the Intermittent compared to the Daily treatment and the Daily treatment compared to the Extreme treatment in the Erken spring experiment.

There were also some lake specific differences in conductivity and pH. Specifically, the runoff treatments had a higher conductivity than the Control in the two Bolmen and the Skogaryd experiments, and pH increased in several or all runoff treatments compared to the Control in the two Erken and the Skogaryd experiments (Table S3), with exception of the Extreme treatment in the Erken spring experiment. In addition, there were a few more differences in pH among runoff treatments, which were not consistent across experiments.

**Text S2. Treatment effects on periphyton growth in the mesocosms**

Periphyton growth in the mesocosms was monitored through the collection of biomass on polyethylene (PE) strips at the end of the experiments (Urrutia Cordero et al. 2021). The biomass was scraped off the

PE strip and resuspended in 50 ml tap water, which was then used for chlorophyll *a* extractions. Treatment, experiment, and their interaction all significantly affected the periphyton biomass that had accumulated on the PE strip in each mesocosm by the end of the experiment (Table S4). Periphyton biomass was significantly higher in the Daily, Extreme and Intermittent treatments compared to the Control (Fig. S18, Table S4) and in the Extreme compared to the Daily treatment. It was also significantly higher in the Bolmen spring experiment compared to all other experiments and higher in the Bolmen summer and Skogaryd compared to the two Erken experiments.

### Supplementary figures

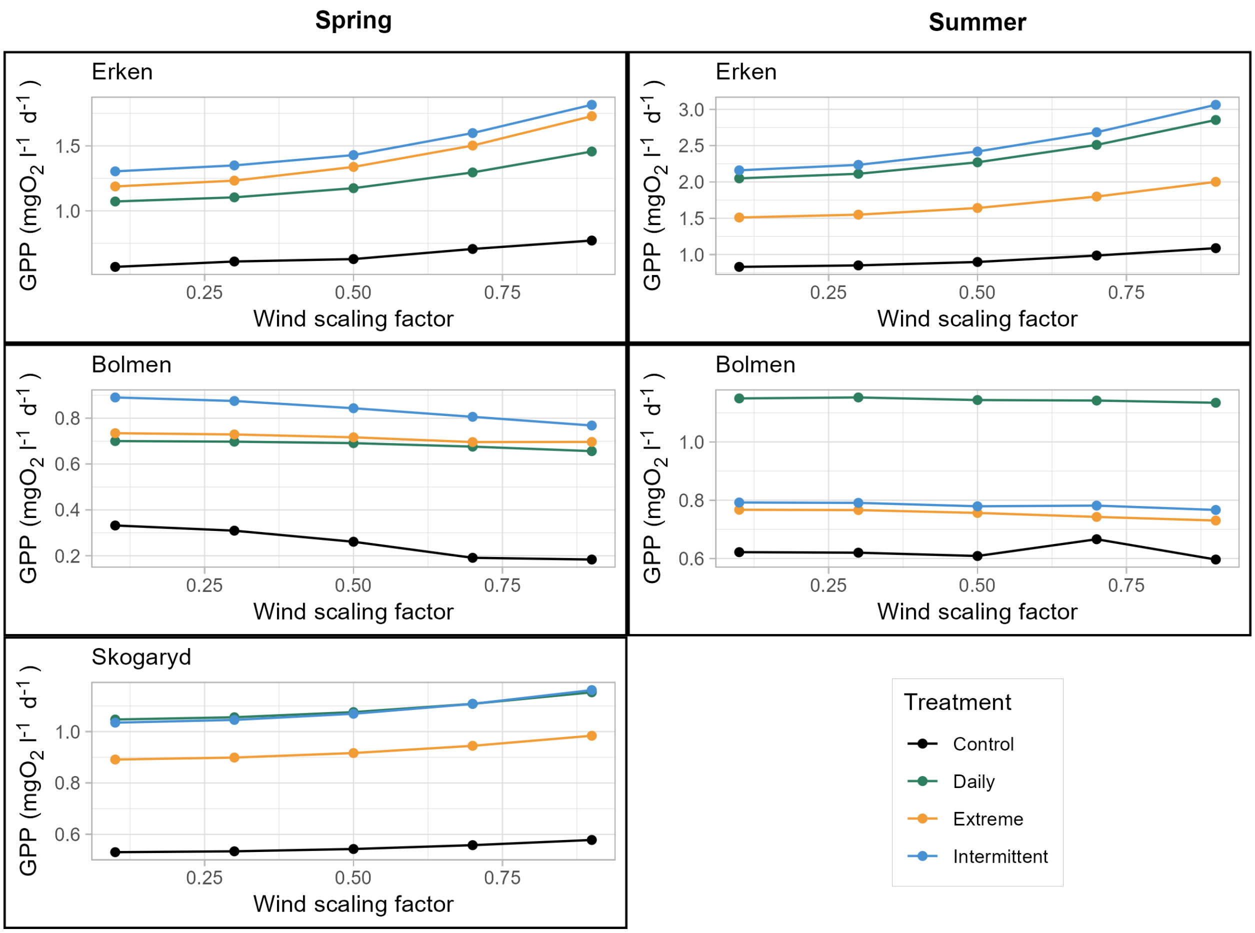

**Figure S1**. Gross Primary production (GPP) over time in the mesocosm experiments using different wind scaling factors (0.1, 0.3, 0.5, 0.7, and 0.9) for the calculations for metabolic rates. The models included in the main manuscript are based on a scaling factor of 0.5.

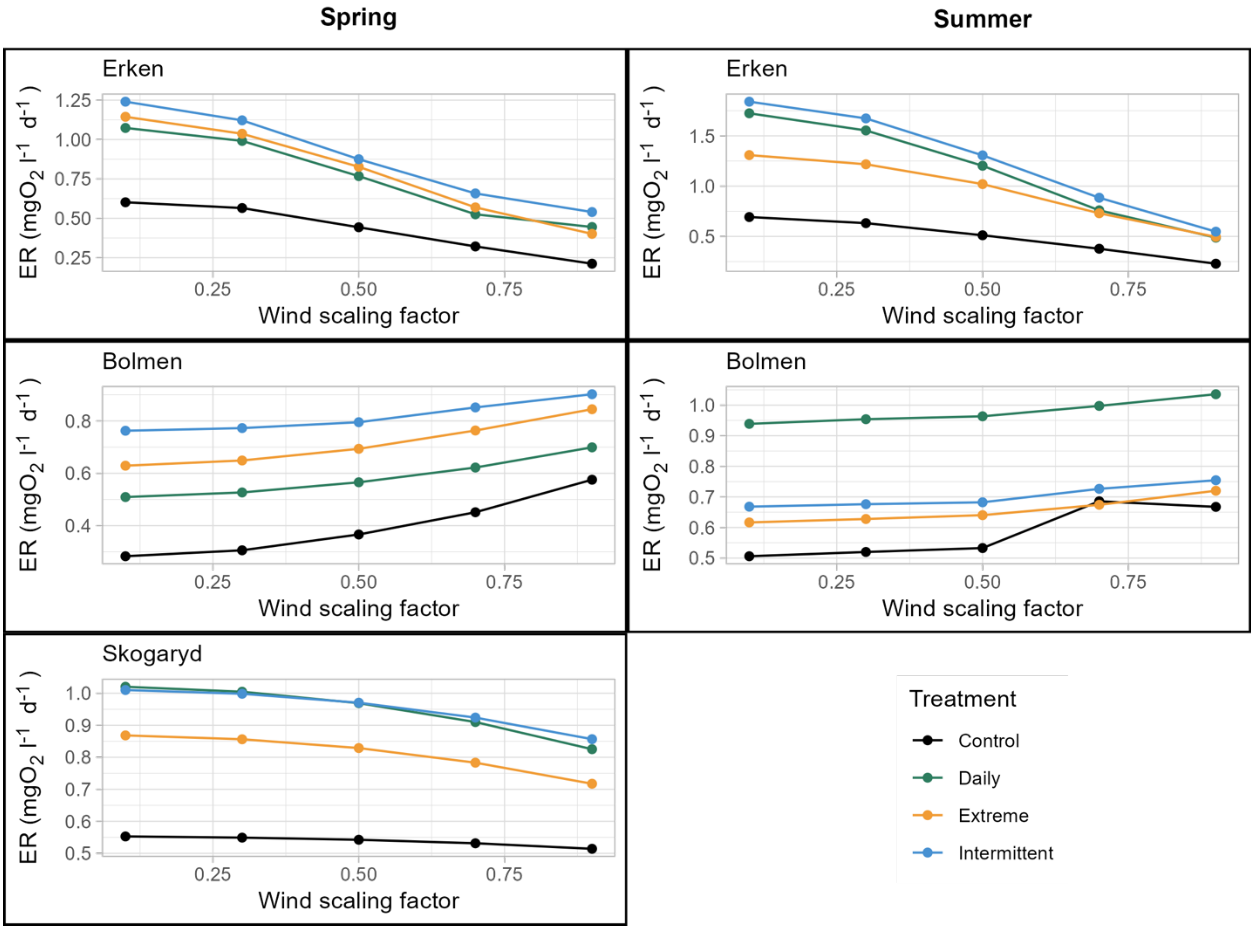

**Figure S2**. Ecosystem respiration (ER) over time in the mesocosm experiments using different wind scaling factors (0.1, 0.3, 0.5, 0.7, and 0.9) for the calculations for metabolic rates. The models included in the main manuscript are based on a scaling factor of 0.5.

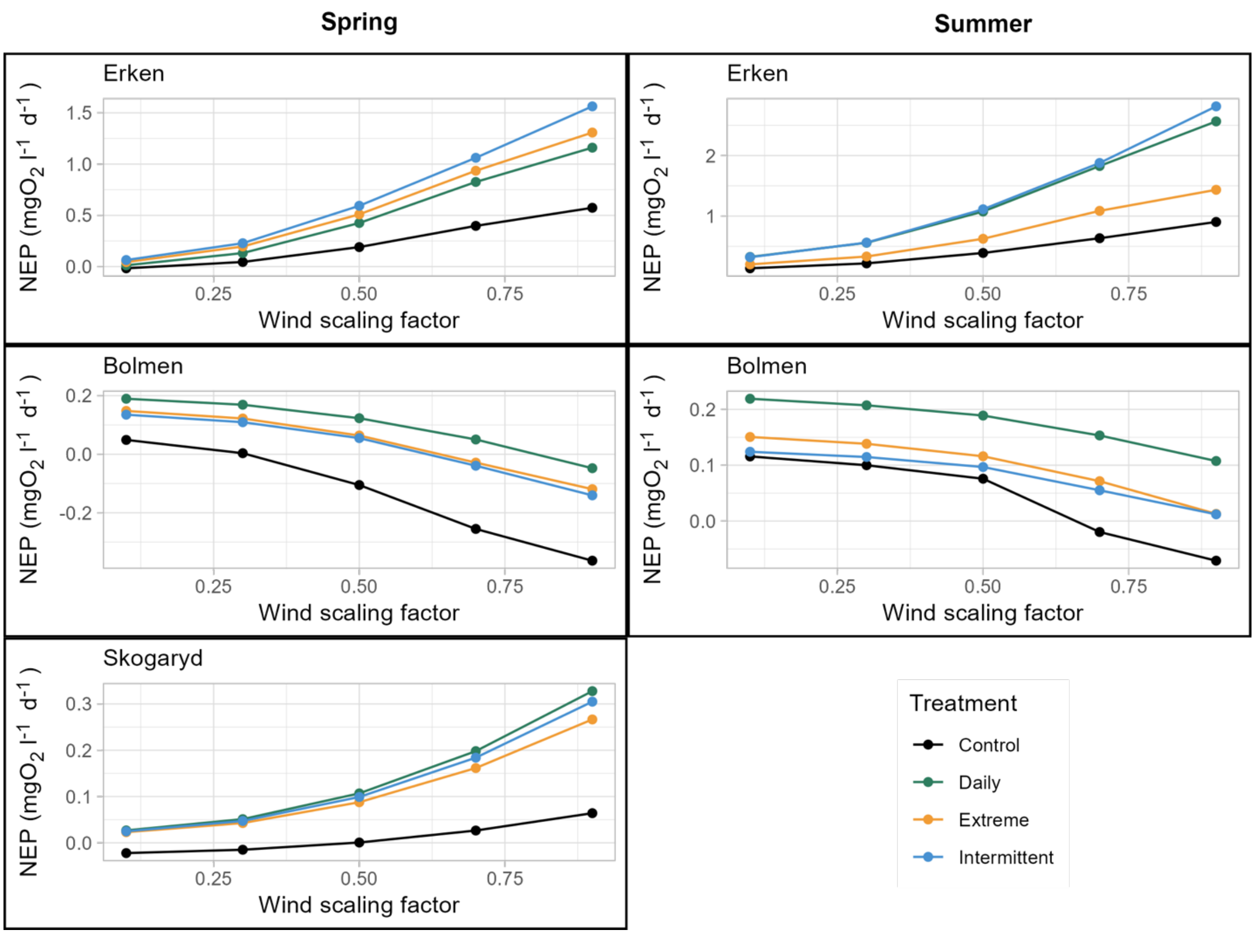

**Figure S3**. Net Ecosystem Production (NEP) over time in the mesocosm experiments using different wind scaling factors (0.1, 0.3, 0.5, 0.7, and 0.9) for the calculations for metabolic rates. The models included in the main manuscript are based on a scaling factor of 0.5

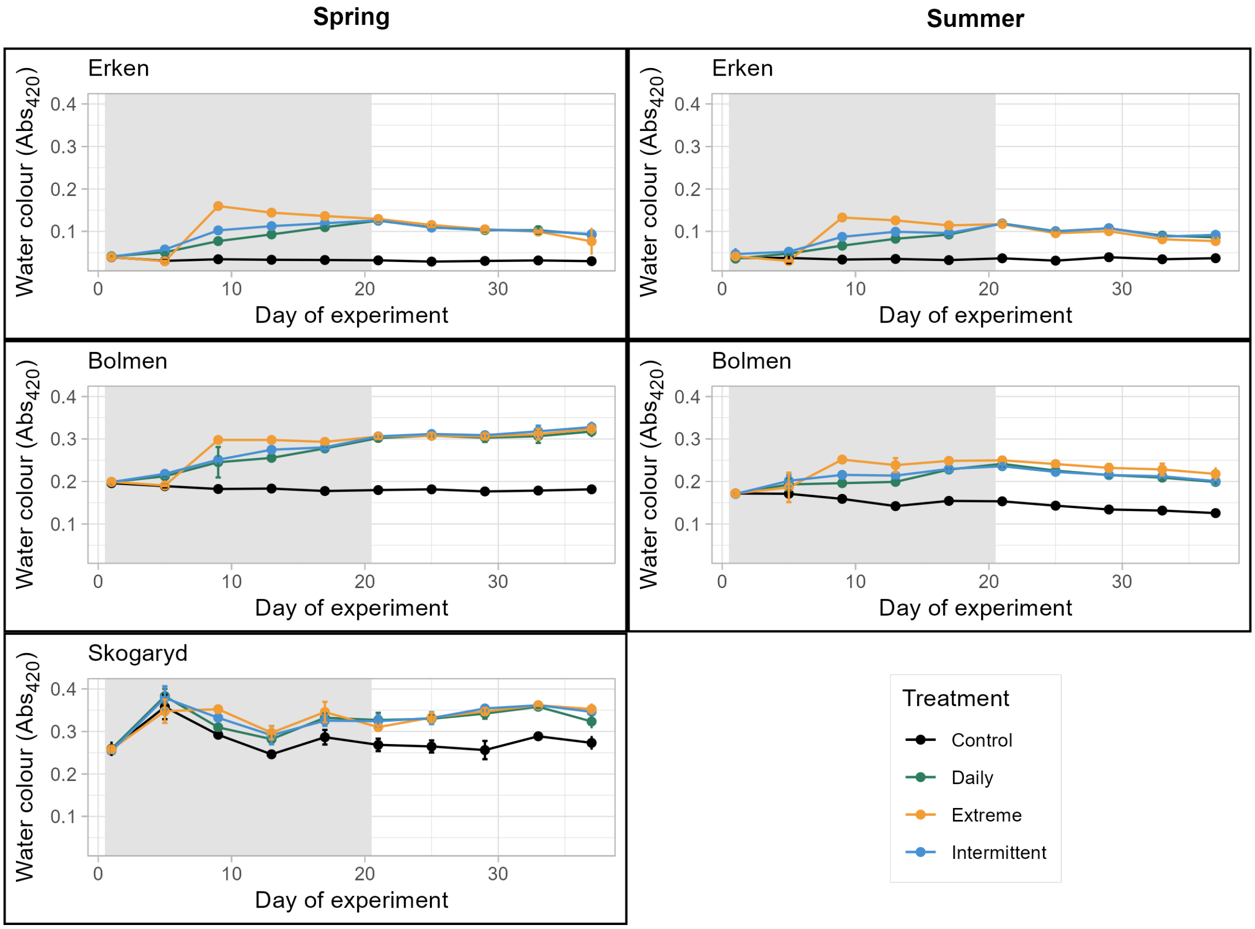

**Figure S4.** Changes in water colour (an estimator of cDOM) measured as absorbance at 420 nm (Abs420) in the mesocosm experiments conducted in spring 2023 and summer 2022 in Erken, Bolmen and Skogaryd. Samples were analyzed every 4th day during the experiments.

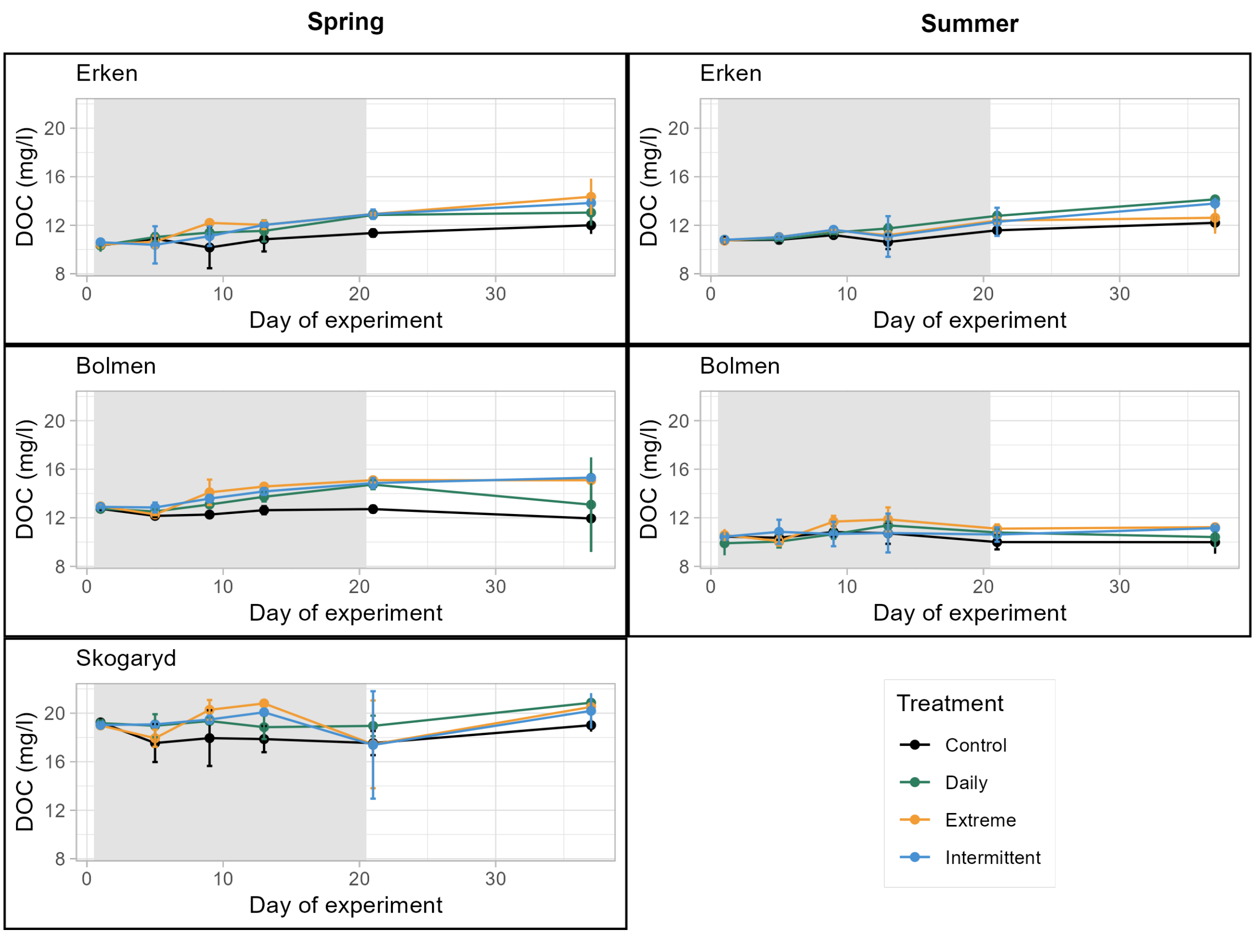

**Figure S5**. Changes in DOC concentrations in the mesocosm experiments conducted in spring 2023 and summer 2022 in Erken, Bolmen and Skogaryd. Samples were analysed on days 1, 5, 9, 13, 21 and 37 of the experiment.

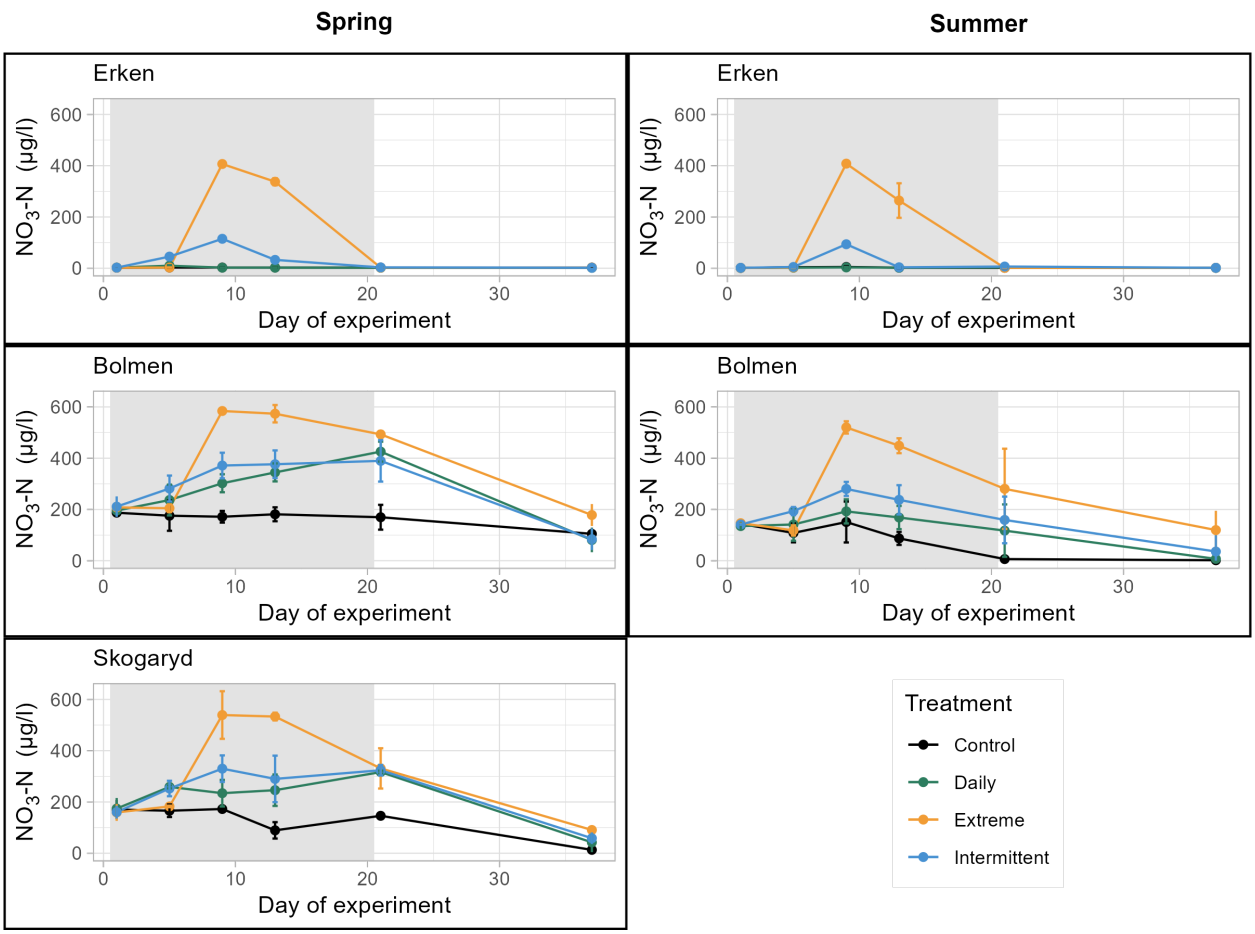

**Figure S6**. Changes in nitrate ($NO_3$-N) concentrations in the mesocosm experiments conducted in spring 2023 and summer 2022 in Erken, Bolmen and Skogaryd. Samples were analysed on days 1, 5, 9, 13, 21 and 37 of the experiment.

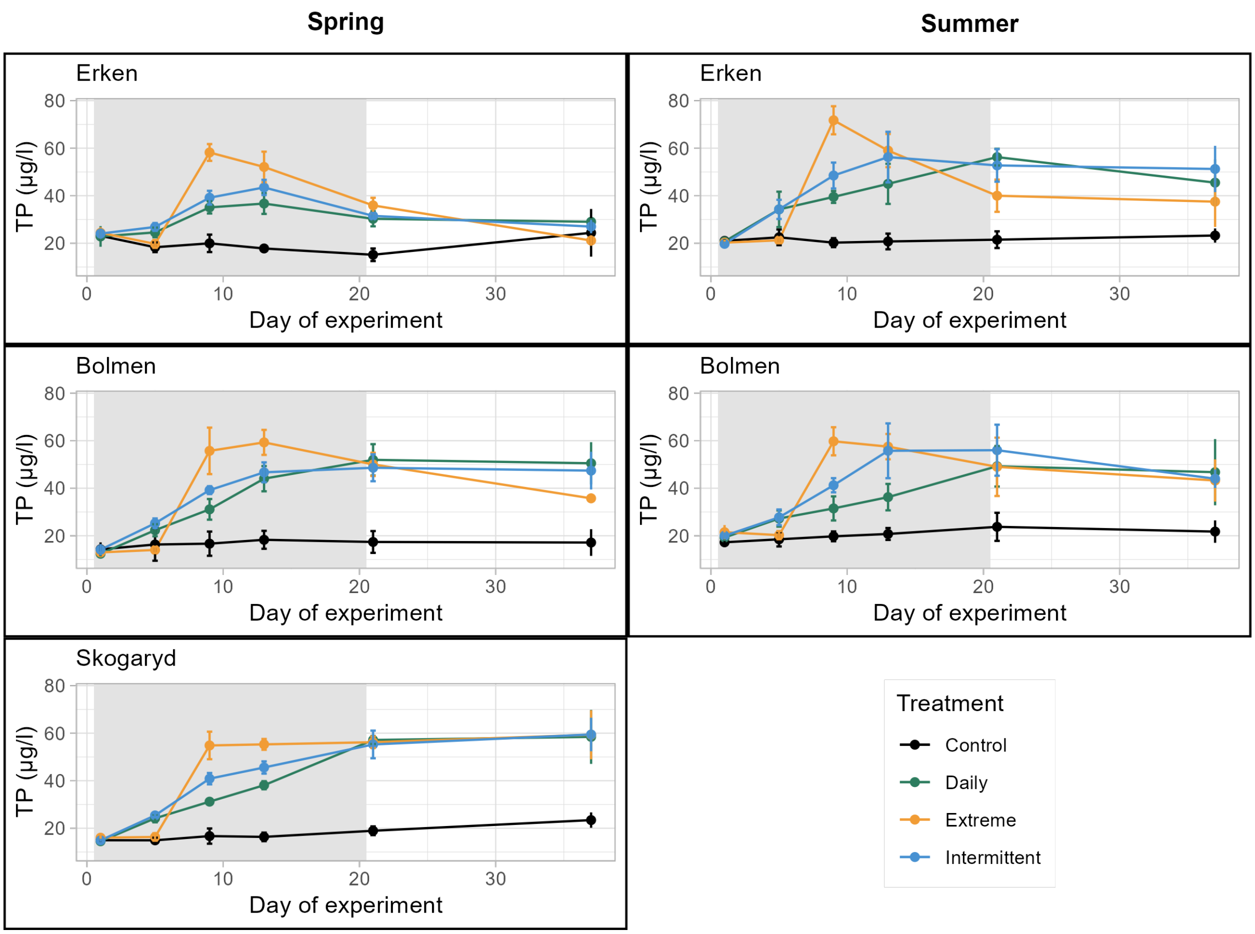

**Figure S7**. Changes in total phosphorus (TP) concentrations in the mesocosm experiments conducted in spring 2023 and summer 2022 in Erken, Bolmen and Skogaryd. Samples were analysed on days 1, 5, 9, 13, 21 and 37 of the experiment.

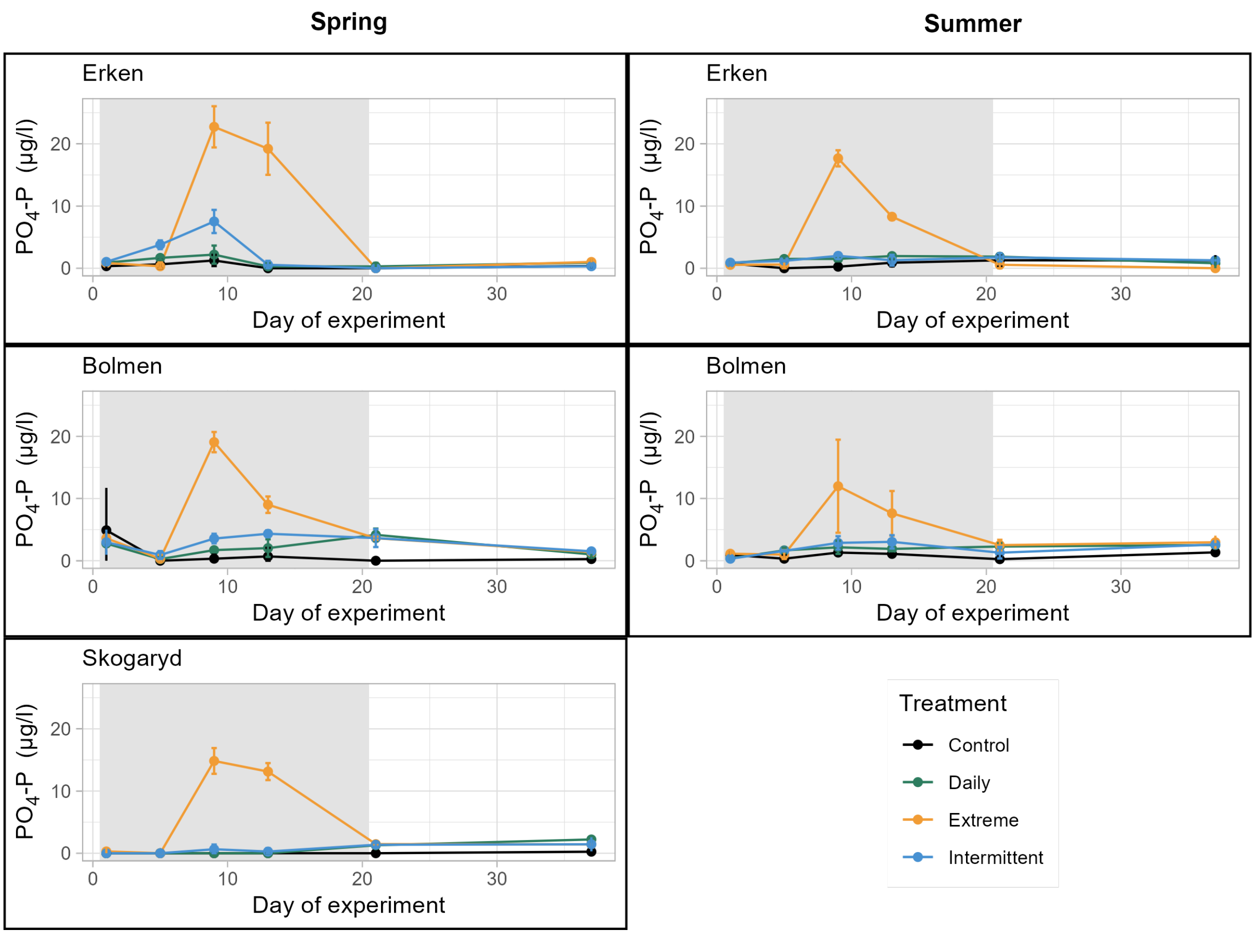

**Figure S8**. Changes in phosphate ($PO_4$-P) concentrations in the mesocosm experiments conducted in spring 2023 and summer 2022 in Erken, Bolmen and Skogaryd. Samples were analysed on days 1, 5, 9, 13, 21 and 37 of the experiment.

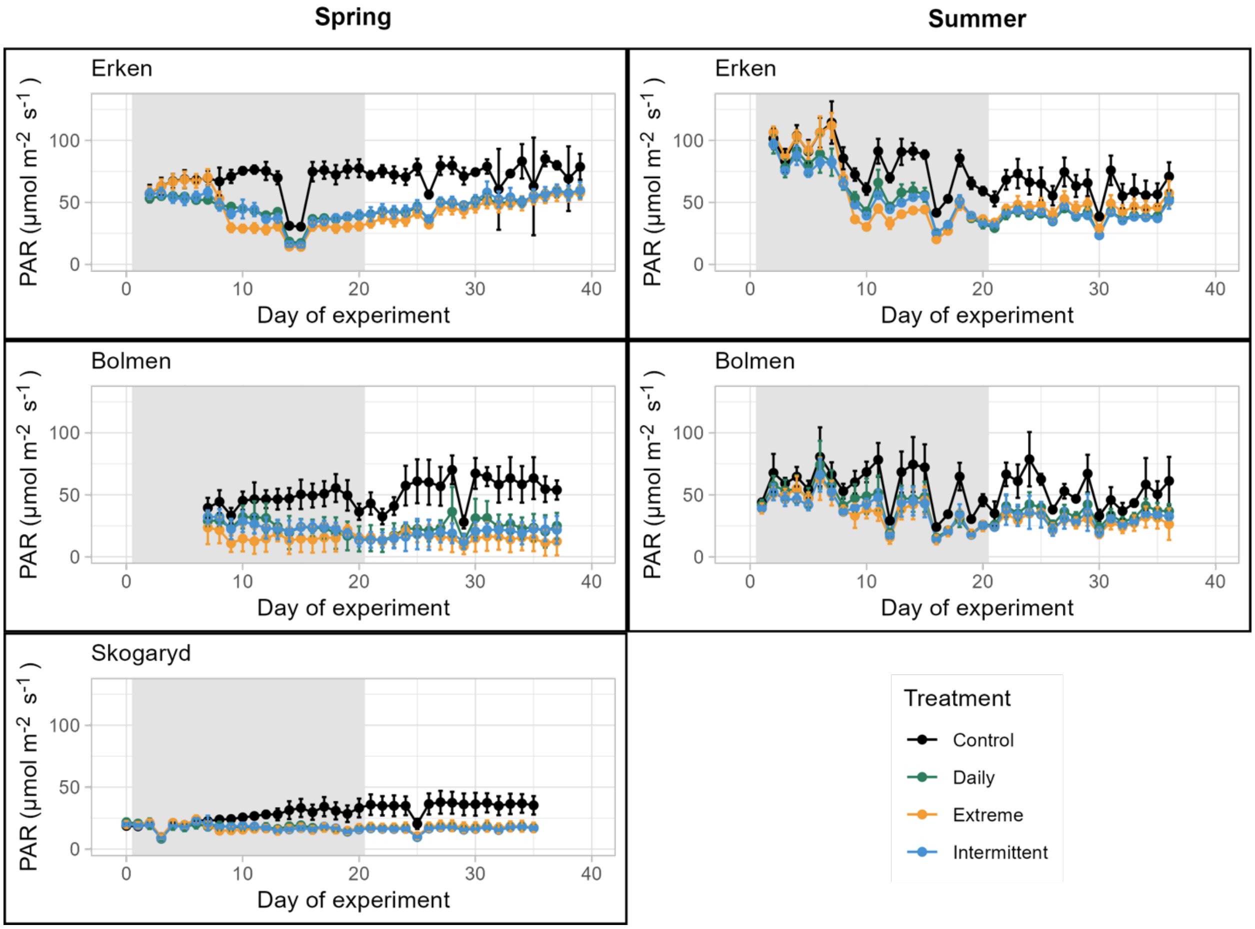

**Figure S9.** Changes in PAR (daily averages) in the mesocosm experiments conducted in spring 2023 and summer 2022 in Erken, Bolmen and Skogaryd.

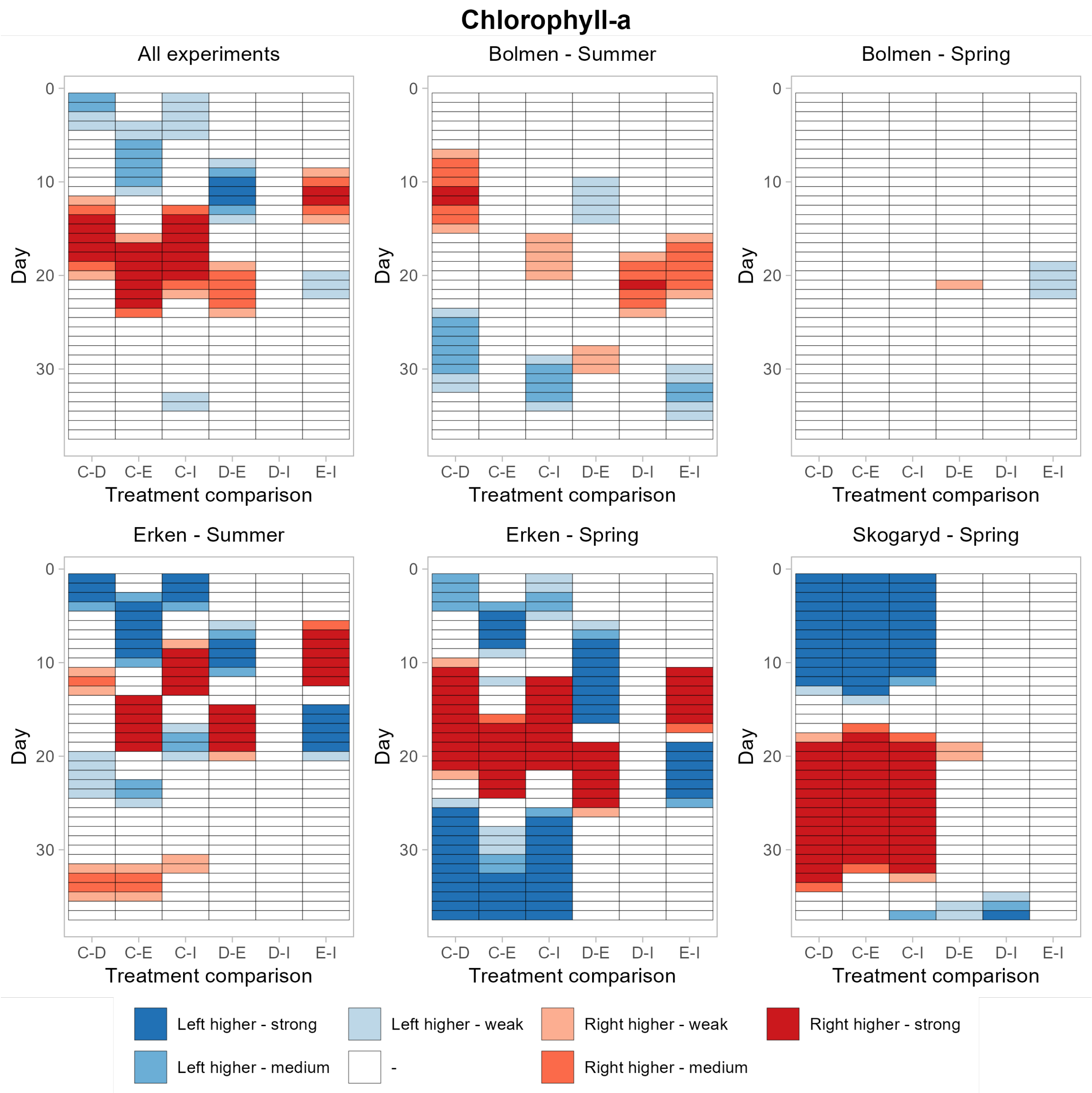

**Figure S10**. Heat maps visualizing significant pairwise differences in chl-*a* concentrations calculated as daily averages from chl-*a* fluorescence between treatments on the different days of the experiment based on GAMM. Colours are split based on *p*-value categories (strong: <0.001, medium: <0.01, weak: <0.05, no effect >0.05.  $R^2$ values for the models were as follows: Bolmen spring: 0.591, Bolmen summer: 0.422, Erken spring: 0.821, Erken summer: 0.532, Skogaryd: 0.909. Abbreviations are as follows: C. Control, D: Daily Treatment, I: Intermittent treatment, E: Extreme treatment. If, for example for pair C-D left is higher, it means that the Control was significantly higher compared to the Daily treatment, whereas 'right higher' means that the Daily treatments was higher compared to the Control. All other comparisons can be read in the same way.

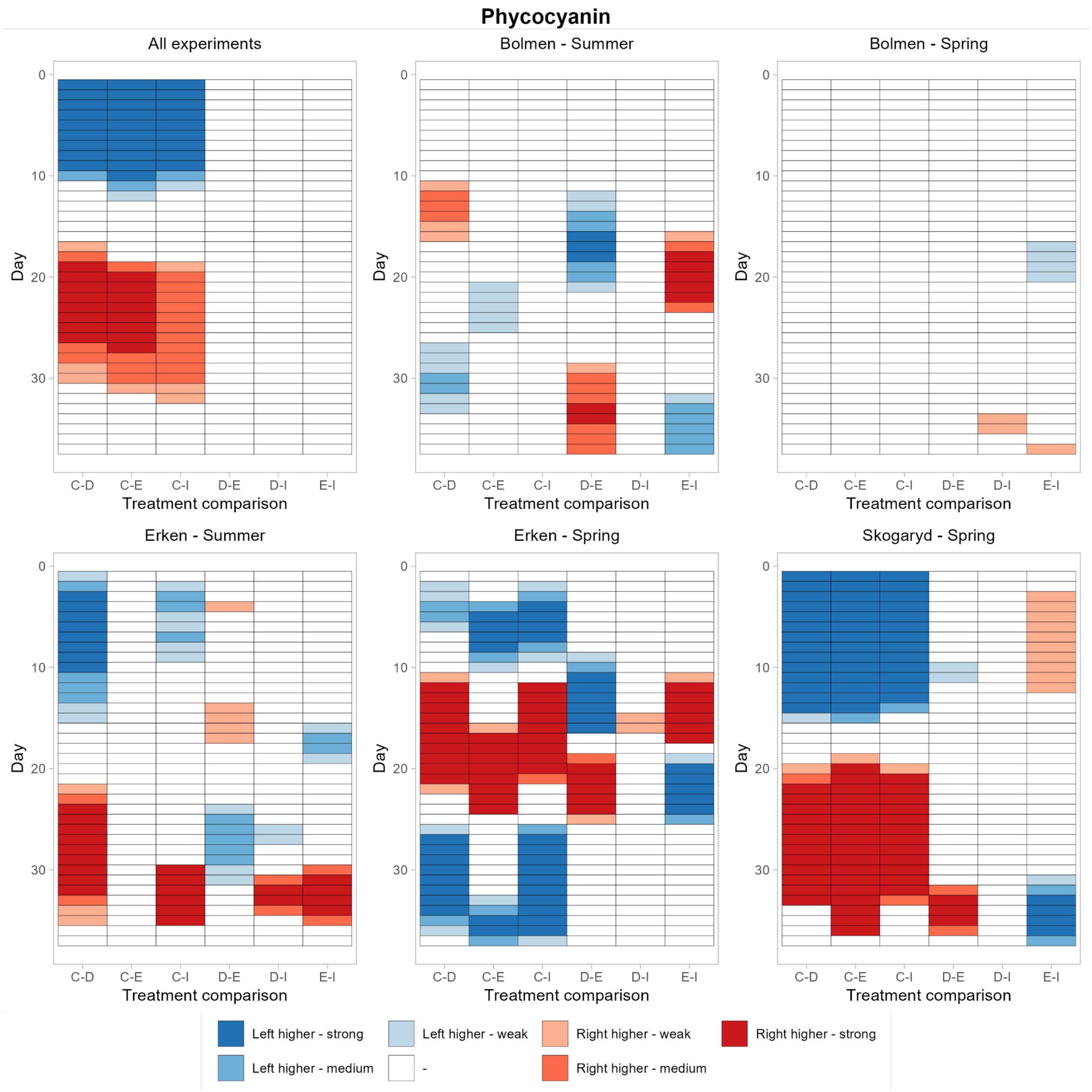

**Figure S11**. Heat maps visualizing significant pairwise differences in phycocyanin concentrations calculated as daily averages from phycocyanin fluorescence between treatments on the different days of the experiment based on GAMM. Colours are split based on *p*-value categories (strong: <0.001, medium: <0.01, weak: <0.05, no effect >0.05. $R^2$ values for the models were as follows: Bolmen spring: 0.263, Bolmen summer: 0.484, Erken spring: 0.717, Erken summer: 0.5, Skogaryd: 0.898. Abbreviations are as follows: C. Control, D: Daily Treatment, I: Intermittent treatment, E: Extreme treatment. If, for example for pair C-D left is higher, it means that the Control was significantly higher compared to the Daily treatment, whereas 'right higher' means that the Daily treatments was higher compared to the Control. All other comparisons can be read in the same way.

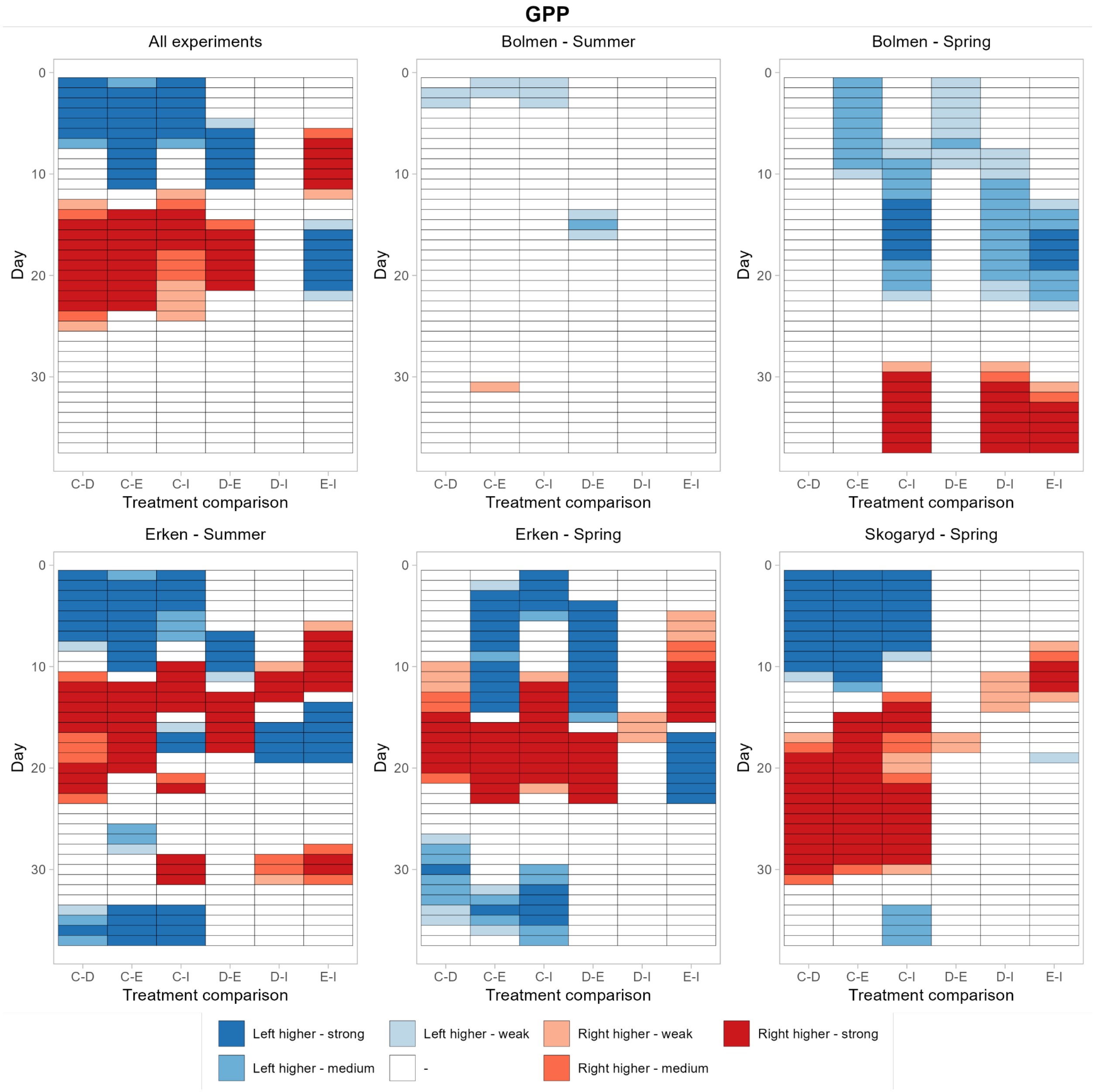

**Figure S12**. Heat maps visualizing significant pairwise differences in GPP between treatments on the different days of the experiment based on GAMM. Colours are split based on *p*-value categories (strong: <0.001, medium: <0.01, weak: <0.05, no effect >0.0. R2 values for the models were as follows: Bolmen spring: 0.691, Bolmen summer: 0.564, Erken spring: 0.615, Erken summer: 0.754, Skogaryd: 0.809. Abbreviations are as follows: C. Control, D: Daily Treatment, I: Intermittent treatment, E: Extreme treatment. If, for example for pair C-D left is higher, it means that the Control was significantly higher compared to the Daily treatment, whereas 'right higher' means that the Daily treatments was higher compared to the Control. All other comparisons can be read in the same way.

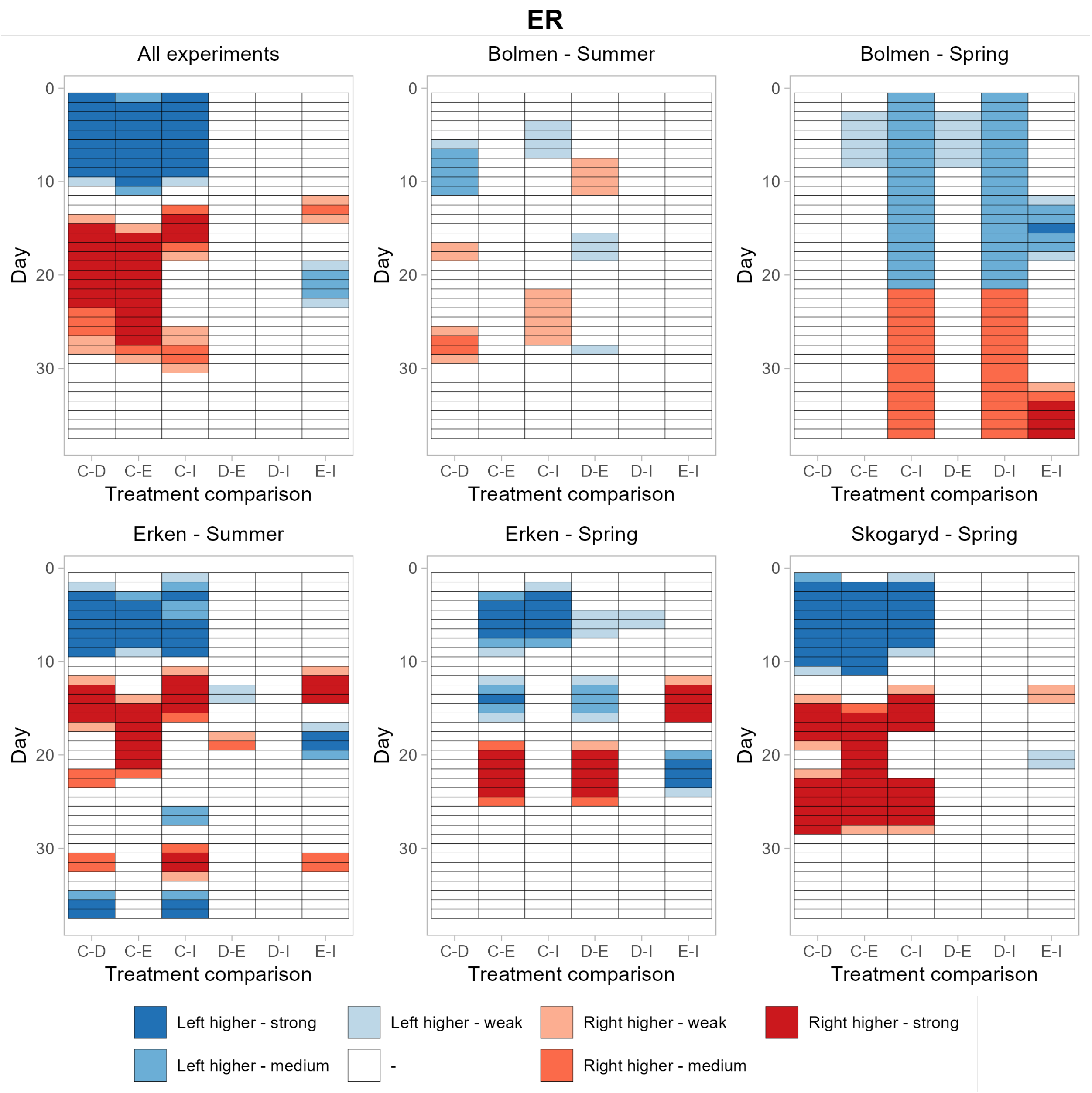

**Figure S13.** Heat maps visualizing significant pairwise differences in ER between treatments on the different days of the experiment based on GAMM. Colours are split based on *p*-value categories (strong: <0.001, medium: <0.01, weak: <0.05, no effect >0.05. $R^2$ values for the models were as follows: Bolmen 2023: 0.241, Bolmen 2022: 0.263, Erken 2023: 0.397, Erken 2022: 0.564, Skogaryd: 0.722. Abbreviations are as follows: C. Control, D: Daily Treatment, I: Intermittent treatment, E: Extreme treatment. If, for example for pair C-D left is higher, it means that the Control was significantly higher compared to the Daily treatment, whereas 'right higher' means that the Daily treatments was higher compared to the Control. All other comparisons can be read in the same way.

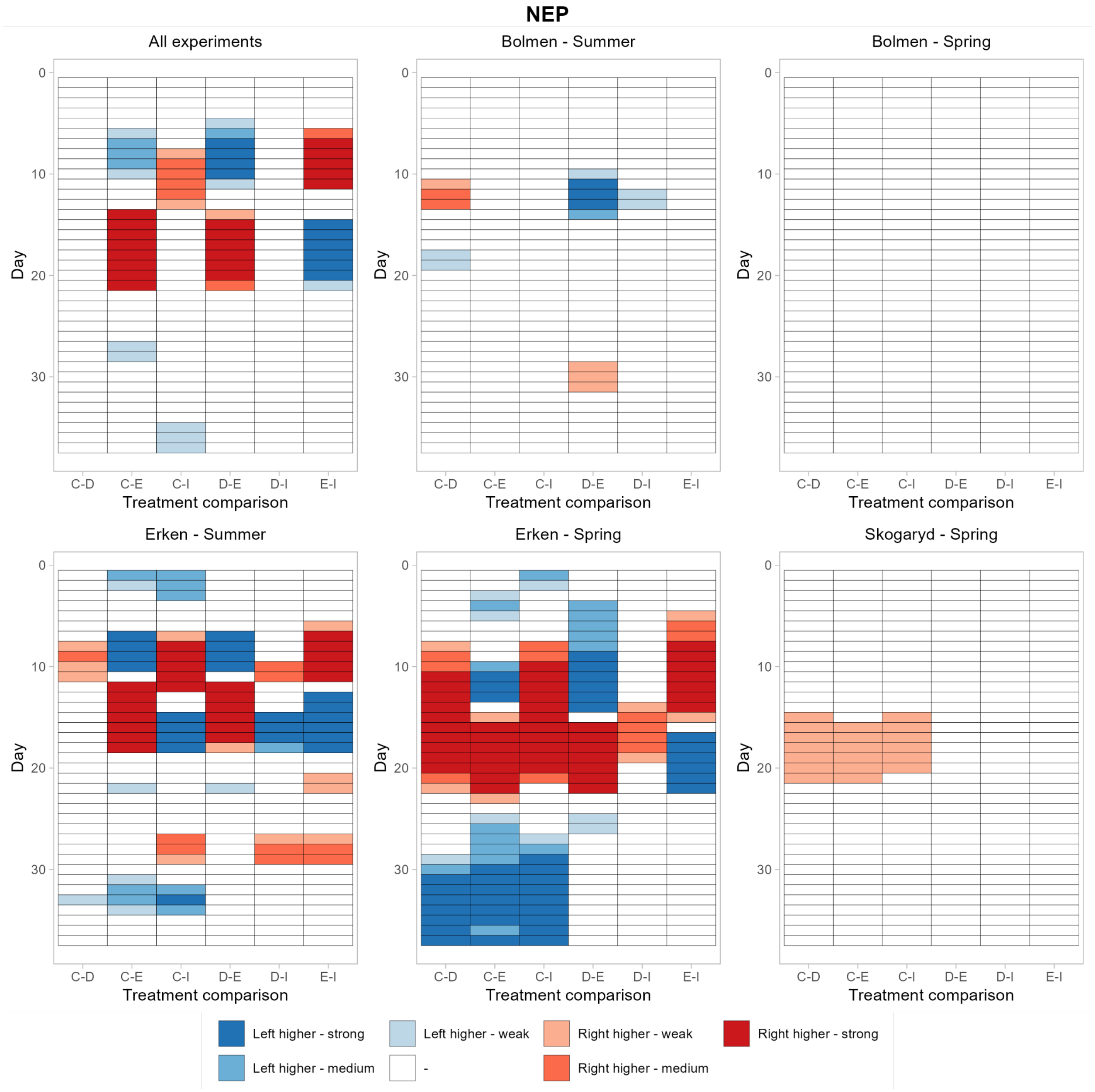

**Figure S14**. Heat maps visualizing significant pairwise differences in NEP between treatments on the different days of the experiment based on GAMM. Colours are split based on *p*-value categories (strong: <0.001, medium: <0.01, weak: <0.05, no effect >0.05. $R^2$ values for the models were as follows: Bolmen 2023: 0.381, Bolmen 2022: 0.167, Erken 2023: 0.622, Erken 2022: 0.446, Skogaryd: 0.073. Abbreviations are as follows: C. Control, D: Daily Treatment, I: Intermittent treatment, E: Extreme treatment. If, for example for pair C-D left is higher, it means that the Control was significantly higher compared to the Daily treatment, whereas 'right higher' means that the Daily treatments was higher compared to the Control. All other comparisons can be read in the same way.

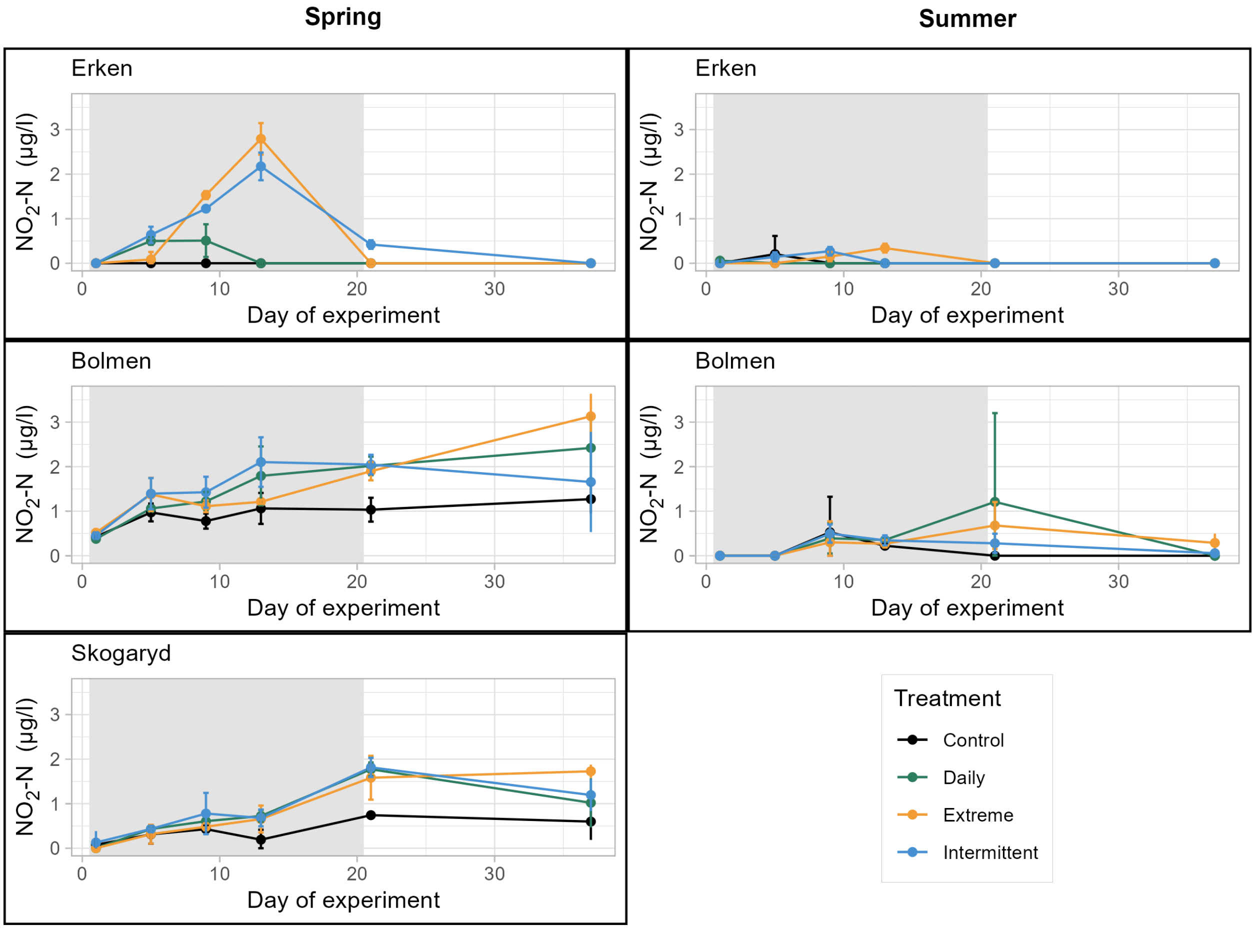

**Figure S15**. Changes in nitrite ($NO_2$-N) concentrations in the mesocosm experiments conducted in spring 2023 and summer 2022 in Lake Erken, Bolmen and Skogaryd. Samples were analysed on days 1, 5, 9, 13, 21 and 37 of the experiment.

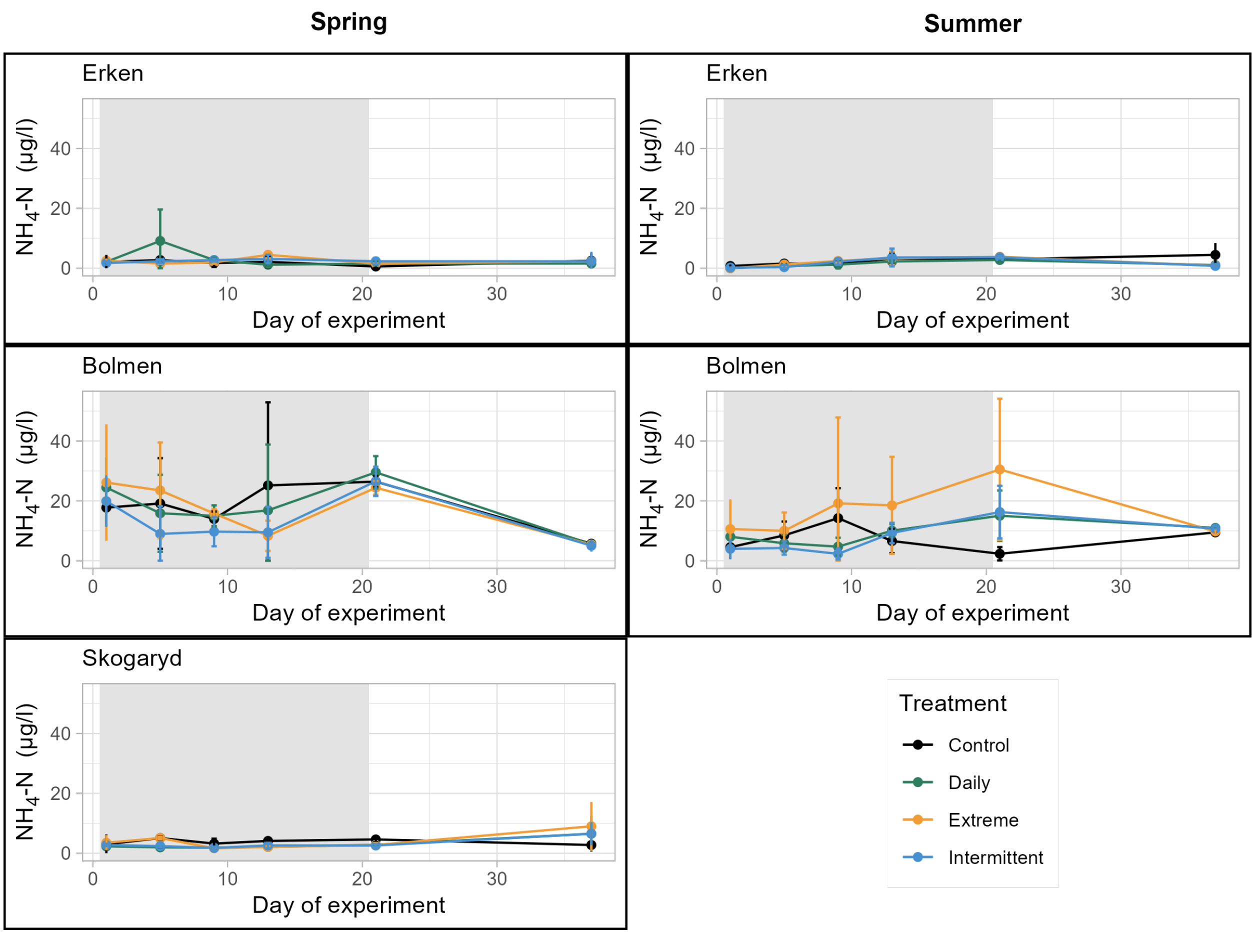

**Figure S16**. Changes in ammonium ($NH_4$-N) concentrations in the mesocosm experiments conducted in spring 2023 and summer 2022 in Lake Erken, Bolmen and Skogaryd. Samples were analysed on days 1, 5, 9, 13, 21 and 37 of the experiment.

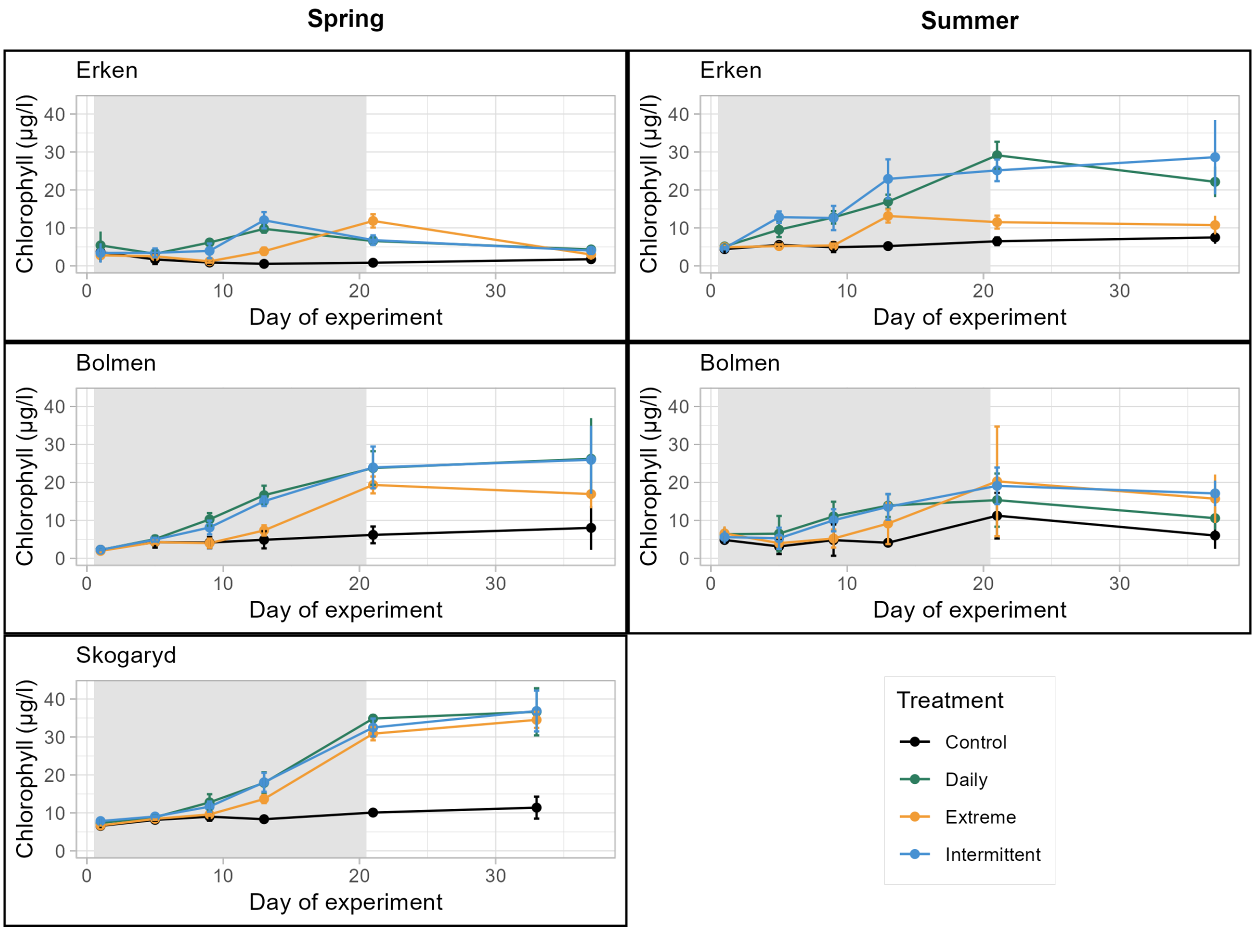

**Figure S17**. Changes in chl-*a* in the mesocosm experiments conducted in spring 2023 and summer 2022 in Lake Erken, Bolmen and Skogaryd. Samples were analysed on days 1, 5, 9, 13, 21 and 37 of the experiments with spectrophotometric analysis of extracted pigments.

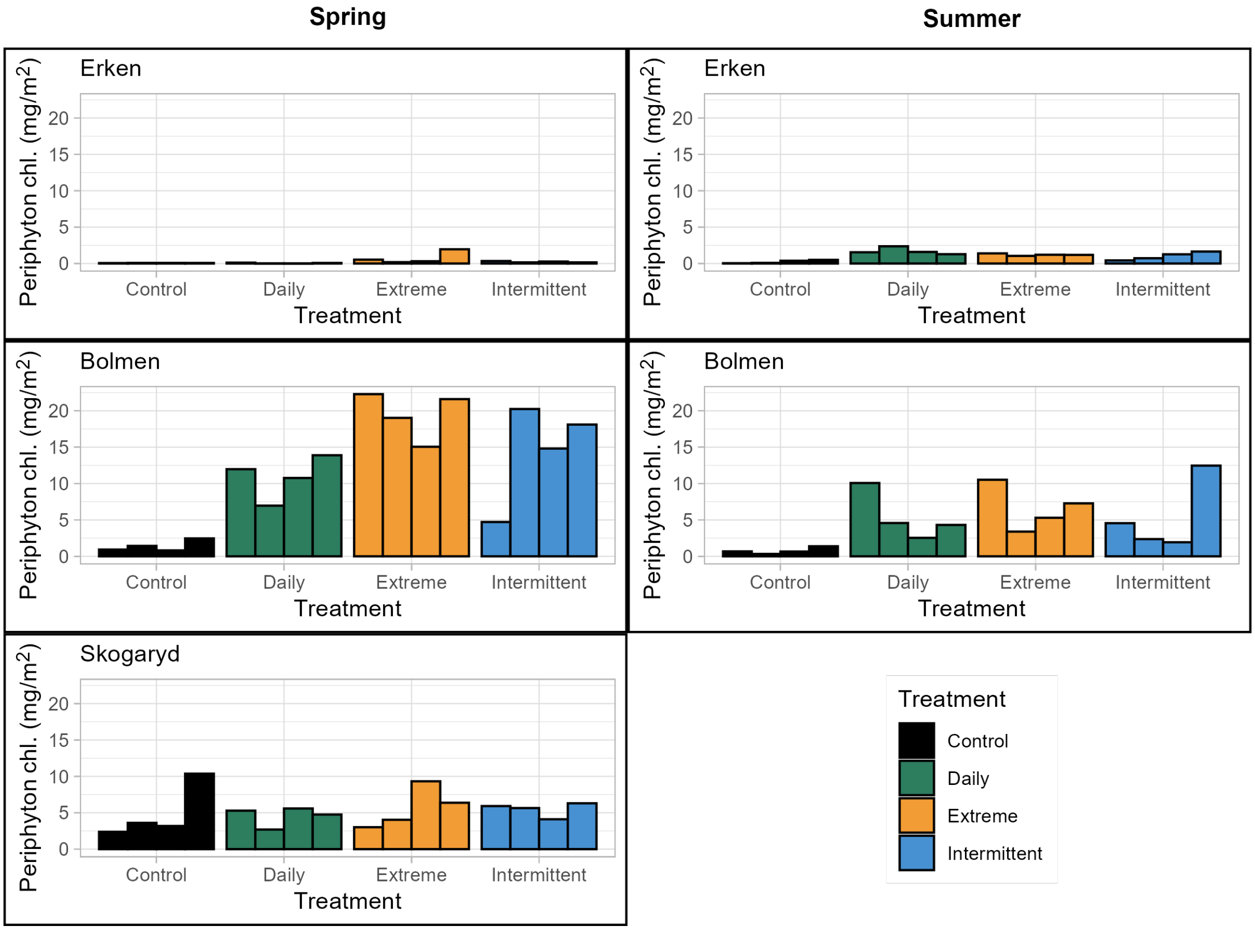

**Figure S18**. Differences in Periphyton biomass estimated as chl-*a* concentration extracted from biofilms growing on polyethylene strips in the mesocosms at the end of the experiments.

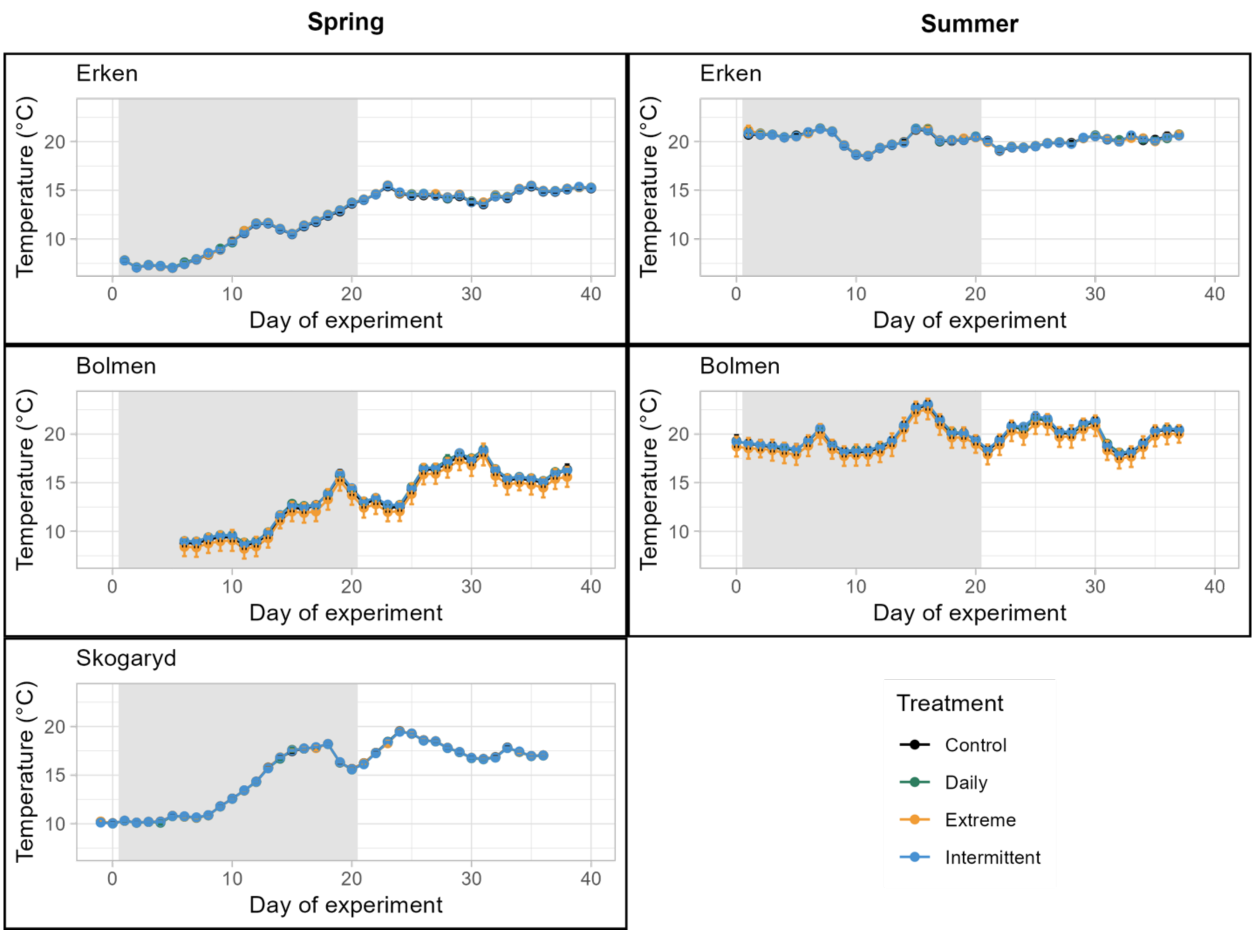

**Fig. S19**. Water temperature (daily averages) in the mesocosms during the experiment conducted in spring 2023 and summer 2022 in Erken, Bolmen and Skogaryd.

### Supplementary Tables

**Table S1.** Environmental conditions in the three lakes during the time periods when the mesocosm experiments were conducted. In all cases, samples and measurements were taken outside the floating platform. Values with ‘<’ indicate that measurements were below the detection limit of the respective method. Average values as well as minimum and maximum values are shown in brackets (). The number of samples (N) is given in the squared bracket. Conductivity (Cond) and pH were measured with a handheld sensor (YSI EXO2 in Bolmen and Erken; hach HQ2220 in Skogaryd) once per day. Water colour (ABS 420) was measured spectrophotometrically every fourth day. Water samples for the determination of chlorophyll *a* concentration by spectrophotometric analysis of extracted chlorophyll *a* as well as concentrations of total phosphorus (TP), dissolved organic carbon (DOC), total dissolved nitrogen (DN), nitrate ($NO_3$-N), nitrite ($NO_2$-N), ammonium ($NH_4$-N) and phosphate ($PO_4$-P) were analysed at selected time point from water samples taken outside of the mesocosm floating platforms. Temperature was measured using an Oxygen Optode 4531 sensor that was mounted in the lake outside of the mesocosm floating platform. Measurements of photosynthetic radiation (PAR) are not from direct measurements in the lake because no consistently measured data were available for all experiments. Instead, the values show daily average of PAR in the control mesocosms throughout the experiment measured using an Appogee SQ-500 sensor mounted at a depth of 40 cm (see methods section and Langenheder et al. 2024 for details).

| | Bolmen summer 2022 | Bolmen spring 2023 | Erken summer 2022 | Erken spring 2023 | Skogaryd spring 2023 |
|---|---|---|---|---|---|
| Cond (µS/cm) | 57.48<br>(54.9 – 61.6)<br>[30] | 60.3<br>(59.4 – 62.6)<br>[35] | 275.27<br>(250.6 – 284.6)<br>[22] | 227.0<br>(181 – 256)<br>[36] | 49.5<br>(47.6 – 51.3)<br>[37] |
| pH | 7.22<br>(7.02 – 7.54)<br>[30] | 6.85<br>(6.65 – 7.7)<br>[36] | 8.52<br>(8.38 – 8.60)<br>[22] | NA | 6.50<br>(5.45 – 7.05)<br>[37] |
| ABS 420 | NA | 0.19<br>(0.195 – 0.206)<br>[10] | NA | 0.037<br>(0.031 -0.04)<br>[10] | 0.301<br>(0.249 – 0.394)<br>[10] |
| Chl-*a* (extracted samples) (µg/L) | 4.83<br>(1.49 – 8.22)<br>[4] | 2.95<br>(2.26 – 4.41)<br>[6] | 11.9<br>(5.77 – 26.8)<br>[4] | 1.23<br>(0.82 – 2.05)<br>[6] | 7.10<br>(5.32 – 9.41)<br>[7] |
| TP (µg/L) | 19.0<br>(16.00– 25.0)<br>[4] | 12.3<br>(10.97 – 14.36)<br>[6] | 21.50<br>(19.0 – 26.0)<br>[4] | 14.87<br>(12.34 – 20.03)<br>[6] | 15.76<br>(11.67 – 21.14)<br>[7] |
| DOC (mg/L) | 9.49<br>(8.29 – 10.36) [4] | 12.44<br>(12.16 – 12.77) [6] | 10.87<br>(10.83 – 10.94) [4] | 10.33<br>(9.09 – 10.84) [6] | 18.24<br>(13.62- 19.79) [7] |

| | | | | | |
|---|---|---|---|---|---|
| DN (µg/L) | 356.12 (275.0 – 418.5) [4] | 624.43 (571.1 – 669.3) [6] | 506.88 (490.9 – 547.7) [4] | 533.02 (449.1 – 605.4) [6] | 573.03 (364.6 – 677.4) [7] |
| $NO_3$-N (µg/L) | 95.0 (56.3 – 112.3) [4] | 223.6 (186.1 – 258.6) [6] | 3.82 (1.66 – 6.54) [4] | 2.81 (1.96 – 5.88) [6] | 122.8 (1.36 – 200.4) [7] |
| $NO_2$-N (µg/L) | 0.23 (<0.15 – 0.48) [4] | 0.64 (0.55 – 0.76) [6] | <0.15 [4] | <0.15 [6] | 0.35 (<0.15 -0.70) [7] |
| $NH_4$-N (µg/L) | 3.07 (<0.78 – 8.41) [4] | 14.74 (3.89 – 27.28) [6] | 1.91 (<0.78 – 2.80) [4] | 2.07 (0.78 – 3.11) [6] | 2.79 (1.75 – 4.38) [7] |
| $PO_4$-P (µg/L) | 1.03 (<0.98 – 1.16) [4] | 1.07 (<0.98 – 1.54) [6] | 1.34 (1.15 – 1.52) [4] | 1.02 (0.98 – 1.21) [6] | <0.98 [7] |
| Temperature (automated sensors) | 19.80 (14.29 – 23.73) [38] | 13.05 (7.33 – 18.67) [33] | 20.40 (19.13 – 21.63) [37] | 12.31 (6.97 – 15.58) [40] | NA |
| PAR (automated sensors) | 54.90 (23.95 - 80.55) [37] | 51.34 (27.94 - 70.10) [33] | 72.63 (38.58 - 114.38) [37] | 68.97 (23.91 - 85.10) [38] | 29.39 (7.82 - 45.41) [38] |

**Table S2.** Results of GAMM analysis across all experiments. The final model included an interaction term between Experiment Day and Treatment with a smoothing factor (s), Enclosure (mesocosm 1-16) nested in Experiment (Bolmen 2022, Bolmen 2023, Erken 2022, Erken 2023 and Skogaryd 2023) as random effects, and Experiment Day in a corAr1 argument to account for the autocorrelation structure in time series data. The global data from the automated sensors (chlorophyll *a* and phycocyanin fluorescence, GPP, ER, NEP and PAR) were too noisy to fit GAMMs with multiple random factors, so Experiment and Enclosure were combined to one random factor (ID). R-sq: adjusted $R^2$ values, i.e. the proportion of variance explained by the model. Edf: effective degrees of freedom.

| Response | R-sq (adj) | Smoothed term | edf | F | p-value |
|---|---|---|---|---|---|
| **Chl- a fluorescence** | 0.454 | s(DAY): C | 1.000 | 0.389 | 0.553 |
| | | s(DAY): D | 4.832 | 7.217 | **<0.001** |
| | | s(DAY): E | 6.525 | 8.657 | **<0.001** |
| | | s(DAY): I | 4.711 | 7.75 | **<0.001** |
| | | s(ID) | 77.000 | 7.758 | **<0.001** |
| **Phycocyanin fluorescence** | 0.52 | s(DAY): C | 1.000 | 0.724 | 0.395 |
| | | s(DAY): D | 3.478 | 22.672 | **<0.001** |
| | | s(DAY): E | 3.200 | 16.394 | **<0.001** |
| | | s(DAY): I | 3.085 | 21.532 | **<0.001** |
| | | s(ID) | 65.238 | 8.094 | **<0.001** |
| **GPP** | 0.639 | s(DAY): C | 1.000 | 29.84 | **<0.001** |
| | | s(DAY): D | 3.665 | 34.56 | **<0.001** |
| | | s(DAY): E | 7.288 | 42.74 | **<0.001** |
| | | s(DAY): I | 4.502 | 34.41 | **<0.001** |
| | | s(ID) | 73.632 | 21.98 | **<0.001** |
| **ER** | 0.384 | s(DAY): C | 1.000 | 20.929 | **<0.001** |
| | | s(DAY): D | 5.102 | 32.165 | **<0.001** |
| | | s(DAY): E | 5.542 | 39.489 | **<0.001** |
| | | s(DAY): I | 7.031 | 31.446 | **<0.001** |
| | | s(ID) | 68.939 | 8.624 | **<0.001** |
| **NEP** | 0.414 | s(DAY): C | 1.000 | 7.122 | **0.0077** |
| | | s(DAY): D | 1.875 | 1.078 | 0.28788 |
| | | s(DAY): E | 6.928 | 12.467 | **<0.001** |
| | | s(DAY): I | 4.227 | 3.691 | **0.00309** |
| | | s(ID) | 70.662 | 11.296 | **<0.001** |
| **Water colour** (Abs420) | 0.961 | s(DAY): C | 1.000 | 16.09 | **<0.001** |

| | | | | | |
|---|---|---|---|---|---|
| | | s(DAY): D | 6.057 | 43.60 | **<0.001** |
| | | s(DAY): E | 8.427 | 50.13 | **<0.001** |
| | | S(DAY): I | 4.899 | 48.97 | **<0.001** |
| | | s(Experiment, Mesocosm) | 78.667 | 239.06 | **<0.001** |
| **DOC** | 0.88 | s(DAY): C | 1.000 | 1.089 | 0.297 |
| | | s(DAY): D | 1.770 | 13.958 | **<0.001** |
| | | s(DAY): E | 3.995 | 13.764 | **<0.001** |
| | | s(DAY): I | 1.000 | 45.166 | **<0.001** |
| | | s(Experiment, Mesocosm) | 77.348 | 42.333 | **<0.001** |
| **$NO_3$-N** | 0.885 | s(DAY): C | 1.000 | 14.10 | **<0.001** |
| | | s(DAY): D | 2.965 | 18.97 | **<0.001** |
| | | s(DAY): E | 4.976 | 219.12 | **<0.001** |
| | | s(DAY): I | 4.357 | 22.55 | **<0.001** |
| | | s(Experiment, Mesocosm) | 76.189 | 27.61 | **<0.001** |
| **Total P** | 0.817 | s(DAY): C | 1.000 | 4.312 | **0.0385** |
| | | s(DAY): D | 3.14 | 91.973 | **<0.001** |
| | | s(DAY): E | 4.964 | 143.572 | **<0.001** |
| | | s(DAY): I | 3.924 | 86.191 | **<0.001** |
| | | s(Experiment, Mesocosm) | 71.936 | 10.333 | **<0.001** |
| **$PO_4$-P** | 0.792 | s(DAY): C | 1.000 | 0.349 | 0.555 |
| | | s(DAY): D | 1.000 | 0.298 | 0.585 |
| | | s(DAY): E | 4.986 | 261.105 | **<0.001** |
| | | s(DAY): I | 2.568 | 3.402 | 0.112 |
| | | s(Experiment, Mesocosm) | 67.999 | 6.271 | **<0.001** |
| **PAR** | 0.76 | s(DAY): C | 8.304 | 13.76 | **<0.001** |
| | | s(DAY): D | 8.191 | 15.51 | **<0.001** |
| | | s(DAY): E | 8.636 | 25.96 | **<0.001** |
| | | s(DAY): I | 8.069 | 13.80 | **<0.001** |
| | | s(ID) | 75.458 | 29.90 | **<0.001** |

| | | | | | |
|---|---|---|---|---|---|
| **$NO_2$-N** | 0.515 | s(DAY): C | 1.000 | 1.120 | **<0.001** |
| | | s(DAY): D | 2.402 | 10.43 | **<0.001** |
| | | s(DAY): E | 2.920 | 14.98 | **<0.001** |
| | | s(DAY): I | 2.979 | 14.78 | **<0.001** |
| | | s(Experiment, Mesocosm) | 65.059 | 4.873 | **<0.001** |
| **$NH_4$-N** | 0.482 | s(DAY): C | 1.000 | 0.444 | 0.5055 |
| | | s(DAY): D | 2.252 | 0.819 | 0.2907 |
| | | s(DAY): E | 2.672 | 2.986 | **0.0201** |
| | | s(DAY): I | 2.830 | 2.232 | **0.0480** |
| | | s(Experiment, Mesocosm) | 65.758 | 5.180 | **<0.001** |
| **Chl-*a* (extracted)** | 0.76 | s(DAY): C | 1.000 | 6.233 | **0.0129** |
| | | s(DAY): D | 4.760 | 58.577 | **<0.001** |
| | | s(DAY): E | 4.779 | 50.846 | **<0.001** |
| | | s(DAY): I | 4.702 | 68.574 | **<0.001** |
| | | s(Experiment, Mesocosm) | 69.538 | 7.309 | **<0.001** |
| **Conductivity** | 0.989 | s(DAY): C | 1.000 | 43.32 | **<0.001** |
| | | s(DAY): D | 1.566 | 26.84 | **<0.001** |
| | | s(DAY): E | 1.621 | 32.42 | **<0.001** |
| | | S(DAY): I | 1.000 | 48.66 | **<0.001** |
| | | S(Experiment, Mesocosm) | 78.903 | 815.63 | **<0.001** |
| **pH** | 0.831 | s(DAY): C | 1.407 | 14.28 | **<0.001** |
| | | s(DAY): D | 3.197 | 35.42 | **<0.001** |
| | | s(DAY): E | 1.955 | 49.05 | **<0.001** |
| | | s(DAY): I | 3.017 | 36.68 | **<0.001** |
| | | s(Experiment, Mesocosm) | 75.322 | 20.75 | **<0.001** |

**Table S3.** T-ratios of pairwise comparisons of interaction terms of treatments and time in GAMMs based on all experiments and separate single experiments. C: Control, D: Daily Treatment, E: Extreme treatment, I: intermittent treatment. Response variables tested were water colour ($ABS_{420}$), concentrations of DOC, total phosphorus (TP), nitrate ($NO_3$-N), nitrite ($NO_2$-N), ammonium (NH4-N) and phosphate

($PO_4$-P), and extracted chlorophyll *a*, as well as pH, Conductivity (Cond), photosynthetically active radiation (PAR) and water temperature.
R-sq: adjusted $R^2$ values, i.e. the proportion of variance explained by the model. Significance codes: *** < 0.001, ** <0.01, * < 0.05, ‘ < 0.1. For units see Table S1.

| | | **All** | **Summer 2022** | **Spring 2023** | **Summer 2022** | **Spring 2023** | **Spring 2023** |
|---|---|---|---|---|---|---|---|
| | | | **Bolmen** | **Bolmen** | **Erken** | **Erken** | **Skogaryd** |
| Water colour ($Abs_{420}$) | R-sq (adj) | 0.961 | 0.938 | 0.979 | 0.98 | 0.92 | 0.877 |
| | C-D | **-6.193*** ** | **-8.536*** ** | **-6.896*** ** | **-14.054*** ** | **-14.039*** ** | -0.921 |
| | C-E | **-5.306*** ** | **-6.7*** ** | **-5.207*** ** | **-14.245*** ** | **-13.808*** ** | 0.235 |
| | C-I | **-4.869*** ** | **-6.943*** ** | **-6.55*** ** | **-11.11*** ** | **-12.939*** ** | 0.567 |
| | D-E | 0.053 | 0.605 | 0.257 | -0.336 | -0.864 | 1.158 |
| | D-I | 1.367 | 1.864 | 0.598 | 1.992 | 0.304 | 1.489 |
| | E-I | 1.202 | 1.087 | 0.243 | 2.306 | 1.132 | 0.333 |
| DOC | R-sq (adj) | 0.88 | 0.174 | 0.474 | 0.732 | 0.682 | 0.254 |
| | C-D | -1.969 | -2.34 | -2.198 | 0 | -0.844 | -0.773 |
| | C-E | **-2.649* ** | -2.03 | -1.759 | 0 | **-3.509** ** | **-2.761* ** |
| | C-I | **-4.014*** ** | 0 | 0 | 0 | **-3.109* ** | -1.072 |
| | D-E | -1.596 | 0.224 | 0.601 | 0 | 0.665 | -2.407 |
| | D-I | 1.595 | 2.34 | 2.198 | 0 | 0.685 | -0.546 |
| | E-I | 2.479 | 2.03 | 1.759 | 0 | 0.685 | 1.839 |
| $NO_3$-N | R-sq (adj) | 0.885 | 0.867 | 0.867 | 0.98 | 0.98 | 0.874 |
| | C-D | **-4.513*** ** | **-2.661* ** | **-6.677*** ** | 0.113 | 0 | **-3.723** ** |
| | C-E | **-10.705** ** | **-5.161*** ** | **-8.402*** ** | **-9.815*** ** | **-54.601*** ** | **-7.572*** ** |
| | C-I | **-2.61* ** | **-3.732** ** | **-7.124*** ** | **5.12*** ** | **9.278*** ** | **-2.628’** |
| | D-E | **-6.982*** ** | **-3.611** ** | **-2.819* ** | **-9.816*** ** | **-54.601*** ** | **-6.25*** ** |
| | D-I | 0.3 | -1.53 | -0.197 | **5.119*** ** | **9.278*** ** | -0.875 |
| | E-I | **6.287*** ** | 2.319 | **2.679* ** | **10.576*** ** | **45.203*** ** | **4.683*** ** |
| Total P | R-sq (adj) | 0.817 | 0.885 | 0.954 | 0.888 | 0.897 | 0.951 |
| | C-D | **-6.258*** ** | **-3.126* ** | **-7.502*** ** | **-4.688*** ** | **-5.39*** ** | **-3.725** ** |
| | C-E | **-7.812*** ** | **-4.72*** ** | **-10.155*** ** | **-3.727** ** | **-7.171*** ** | **-4.884*** ** |
| | C-I | **-8.797*** ** | **-7.455*** ** | **-7.732*** ** | **-6.067*** ** | **-7.131*** ** | **-5.936*** ** |

| | | | | | | | |
|---|---|---|---|---|---|---|---|
| | D-E | **-3.342**** | **-2.755*** | **-4.271***** | -0.846 | -2.528 | -2.05 |
| | D-I | **-3.066*** | **-4.235***** | -0.608 | -1.63 | -1.955 | -1.435 |
| | E-I | 0.666 | -0.534 | **3.671**** | -0.414 | 0.704 | 0.97 |
| $PO_4$-P | R-sq (adj) | 0.792 | 0.663 | 0.832 | 0.97 | 0.947 | 0.98 |
| | C-D | -0.804 | 0 | -1.899 | **-2.77*** | 0 | -0.553 |
| | C-E | **-6.162***** | -1.514 | -0.149 | -0.008 | **-11.352***** | **-18.78***** |
| | C-I | -1.871 | 0 | -2.368 | -0.998 | **4.336***** | -2.469 |
| | D-E | **-6.136***** | -1.514 | 1.175 | 1.103 | **-11.352***** | **-17.23***** |
| | D-I | -1.821 | 0 | -0.219 | 1.912 | **4.336***** | -1.059 |
| | E-I | **4.663***** | 1.514 | -1.38 | -0.234 | **11.171***** | **17.95***** |
| PAR | R-sq (adj) | 0.76 | 0.541 | 0.822 | 0.659 | 0.832 | 0.787 |
| | C-D | 1.6040 | 0.397 | -0.866 | 1.414 | **5.616***** | **4.431***** |
| | C-E | 0.9670 | -1.438 | **-4.503***** | 1.489 | **7.275***** | **4.703***** |
| | C-I | 0.6220 | -1.232 | -1.794 | 0.551 | **5.321***** | **4.131***** |
| | D-E | -0.6290 | -2.119 | **-4.826***** | 0.085 | 1.893 | 0.379 |
| | D-I | -1.0030 | -1.878 | -1.117 | -0.873 | -0.239 | 0.162 |
| | E-I | -0.3600 | -1.878 | **3.653**** | -0.952 | -2.102 | -0.171 |
| $NO_2$-N | R-sq (adj) | 0.515 | 0.13 | 0.696 | 0.371 | 0.971 | 0.84 |
| | C-D | **-2.821*** | -2.331 | -2.311 | 0 | **3.704**** | -1.969 |
| | C-E | **-4.164***** | 0 | 0 | **-4.871***** | **-29.14***** | -2.528 |
| | C-I | **-4.972***** | 0 | **-4.144***** | 1.342 | **-21.188***** | 0.211 |
| | D-E | -1.366 | 2.331 | 2.311 | **-4.871***** | **-23.836***** | -0.134 |
| | D-I | -2.039 | 2.331 | -1.844 | 1.342 | **-17.964***** | 1.436 |
| | E-I | -0.639 | 0 | **-4.144***** | **4.428***** | **5.79***** | 1.635 |
| $NH_4$-N | R-sq (adj) | 0.482 | 0.557 | 0.143 | 0.51 | 0.209 | 0.494 |
| | C-D | -0.919 | 1.224 | 1.586 | **-2.629*** | 1.883 | 0.319 |
| | C-E | -1.25 | **-3.202*** | 1.586 | **-4.175***** | 0 | 2.079 |
| | C-I | -1.006 | 1.808 | 0.649 | **-4.371***** | 0 | 1.07 |
| | D-E | -0.341 | **-3.231**** | 0.001 | -1.483 | -1.883 | 1.957 |
| | D-I | -0.192 | 0.606 | -0.287 | -1.697 | -1.883 | 0.724 |
| | E-I | 0.131 | **3.245**** | -0.287 | -0.221 | 0 | -1.701 |
| Chl-*a* (extracted) | R-sq (adj) | 0.76 | 0.564 | 0.861 | 0.853 | 0.806 | 0.963 |

| | | | | | | | |
|---|---|---|---|---|---|---|---|
| | C-D | **-3.306**** | -2.61 | **-3.635**** | **-3.237**** | **-5.985***** | -0.016 |
| | C-E | -0.356 | -0.741 | -0.676 | -2.343 | -2.437 | 2.018 |
| | C-I | **-3.658**** | -2.233 | **-2.905*** | **-4.198***** | **-9.59***** | -0.136 |
| | D-E | 2.083 | 0.87 | 1.697 | 1.505 | **2.89*** | 1.444 |
| | D-I | -0.233 | 0.276 | 0.367 | -0.014 | **-3.268**** | -0.082 |
| | E-I | -2.323 | -0.651 | -1.333 | -1.8 | **-6.066***** | -1.566 |
| pH | R-sq (adj) | 0.831 | 0.673 | 0.896 | 0.984 | 0.192 | 0.899 |
| | C-D | **-2.883*** | 1.234 | 1.926 | **-5.805***** | **-3.401**** | **-4.959***** |
| | C-E | -0.616 | 1.913 | 1.629 | **-15.346***** | 2.497 | **-3.401**** |
| | C-I | -2.548 | 0.968 | **2.837*** | **-3.308**** | **-3.67**** | **-3.852***** |
| | D-E | 2.185 | 0.665 | 0.075 | **-9.858***** | **3.735**** | 1.921 |
| | D-I | 0.327 | -0.271 | 0.84 | 2.265 | -0.273 | 1.321 |
| | E-I | -1.858 | -0.942 | 0.651 | **11.664***** | **-3.994***** | -0.588 |
| Cond | R-sq (adj) | 0.989 | 0.868 | 0.963 | 0.0755 | 0.82 | 0.963 |
| | C-D | -1.387 | **-3.938***** | **-5.582***** | -0.001 | 0.032 | **-6.987***** |
| | C-E | -1.341 | **-3.259**** | **-8.009***** | 0 | 1.408 | **-7.274***** |
| | C-I | 0.277 | **-2.844*** | **-6.779***** | 0 | 0.882 | **-6.372***** |
| | D-E | -0.009 | 0.471 | **-3.076*** | 0.001 | 1.477 | -0.918 |
| | D-I | 1.396 | 0.761 | -1.305 | 0 | 0.913 | 0.449 |
| | E-I | 1.35 | 0.29 | 1.855 | 0 | 0.516 | 1.312 |

**Table S4.** Result from Tukey HSD post hoc test from an ANOVA testing difference in periphyton growth with treatment and experiment as individual factors and as an interaction terms. There were significant effects of treatment (F-value: 14.512, $p<0.001$), Experiment (F-value: 46.972, $p> 0.001$) and Treatment x Experiment (F-value: 5.991, $P< 0.001$) on periphyton biomass at the end of the mesocosm experiment in Erken (summer 2022, spring 2023), Bolmen (summer 2022, spring 2023) and Skogaryd (spring 2023). D: Daily Addition treatment, E: Extreme addition treatment, I: Intermittent addition treatment, C: Control treatment. Tukey multiple comparisons of means from global treatments and experiment specific treatments were computed with the R function "TukeyHSD".

| | **diff** | **$p_{adj}$** |
|---|---|---|
| **TREATMENT** | | |
| D-C | 0.862 | **0.003** |
| E-C | 3.091 | **<0.001** |
| I-C | 1.648 | **<0.001** |
| E-D | 0.045 | **0.044** |
| I-D | -1.398 | 0.777 |
| I-E | -3.626 | 0.309 |
| **EXPERIMENT** | | |
| Bolmen spring - Bolmen summer | 7.045 | **<0.001** |
| Erken summer -Bolmen summer | -3.480 | **0.003** |
| Erken spring - Bolmen summer | -4.244 | **<0.001** |
| Skogaryd spring - Bolmen summer | 0.639 | 0.958 |
| Erken summer - Bolmen spring | -10.525 | **<0.001** |
| Erken spring - Bolmen spring | -11.289 | **<0.001** |
| Skogaryd spring - Bolmen spring | -6.406 | **<0.001** |
| Erken spring - Erken spring | -0.765 | 0.921 |
| Skogaryd spring - Erken summer | 4.119 | **<0.001** |
| Skogaryd spring - Erken spring | 4.883 | **<0.001** |
